# Sequential unfolding mechanisms of monomeric caspases

**DOI:** 10.1101/2023.01.04.522771

**Authors:** Isha Joglekar, A. Clay Clark

## Abstract

Caspases are evolutionarily conserved cysteinyl proteases that are integral in cell development and apoptosis. All apoptotic caspases evolved from a common ancestor into two distinct subfamilies with either monomeric (initiators) or dimeric (effectors) oligomeric states. The regulation of apoptosis is influenced by the activation mechanism of the two subfamilies, but the evolution of the well-conserved caspase-hemoglobinase fold into the two subfamilies is not well understood. We examined the folding landscape of monomeric caspases from two coral species over a broad pH range of 3 to 10.5. On an evolutionary timescale, the two coral caspases diverged from each other approximately 300 million years ago, and they diverged from human caspases about 600 million years ago. Our results indicate that both proteins have overall high stability, ∼ 15 kcal mol^-1^ near the physiological pH range (pH 6 to pH 8), and unfold via two partially folded intermediates, I_1_ and I_2_, that are in equilibrium with the native and the unfolded state. Like the dimeric caspases, the monomeric coral caspases undergo a pH-dependent conformational change resulting from the titration of an evolutionarily conserved site. Data from molecular dynamics simulations paired with limited proteolysis and MALDI-TOF mass spectrometry show that the small subunit of the monomeric caspases is unstable and unfolds prior to the large subunit. Overall, the data suggest that all caspases share a conserved folding landscape, that a conserved allosteric site can be fine-tuned for species-specific regulation, and that the subfamily of stable dimers may have evolved to stabilize the small subunit.

## Introduction

Caspases are a family of cysteine proteases and are well known for their role in apoptosis and inflammation^1^. At lower activation levels, however, caspases play essential roles in non-apoptotic functions like cell proliferation and differentiation, tissue regeneration, and neuronal development.^1,2^ All caspases exist as inactive zymogens in the cell, and the caspase protomer comprises an N-terminal pro-domain and a large and a small subunit that are connected by an inter-subunit linker ^3^ (Fig. 1A). The apoptotic caspases evolved into two subfamilies and are classified as either initiator or effector caspases, depending on their entry into the cell death cascade. Initiator caspases exist as monomers and require dimerization for complete activation, whereas the effector caspases are dimers that require processing for complete activation. ^4–6^ Within the protomer of monomeric caspases, the pro-domain consists of an N-terminal CARD (caspase activation and recruitment domain) or two DEDs (death effector domain), which link caspase activation to death activation platforms in the cell, such as the death inducing signaling complex (DISC). In contrast, the dimeric effector caspases consist of short pro-domains and are activated via cleavage of the intersubunit linker by the initiator caspases. ^1,7–9^

**Figure 1.**
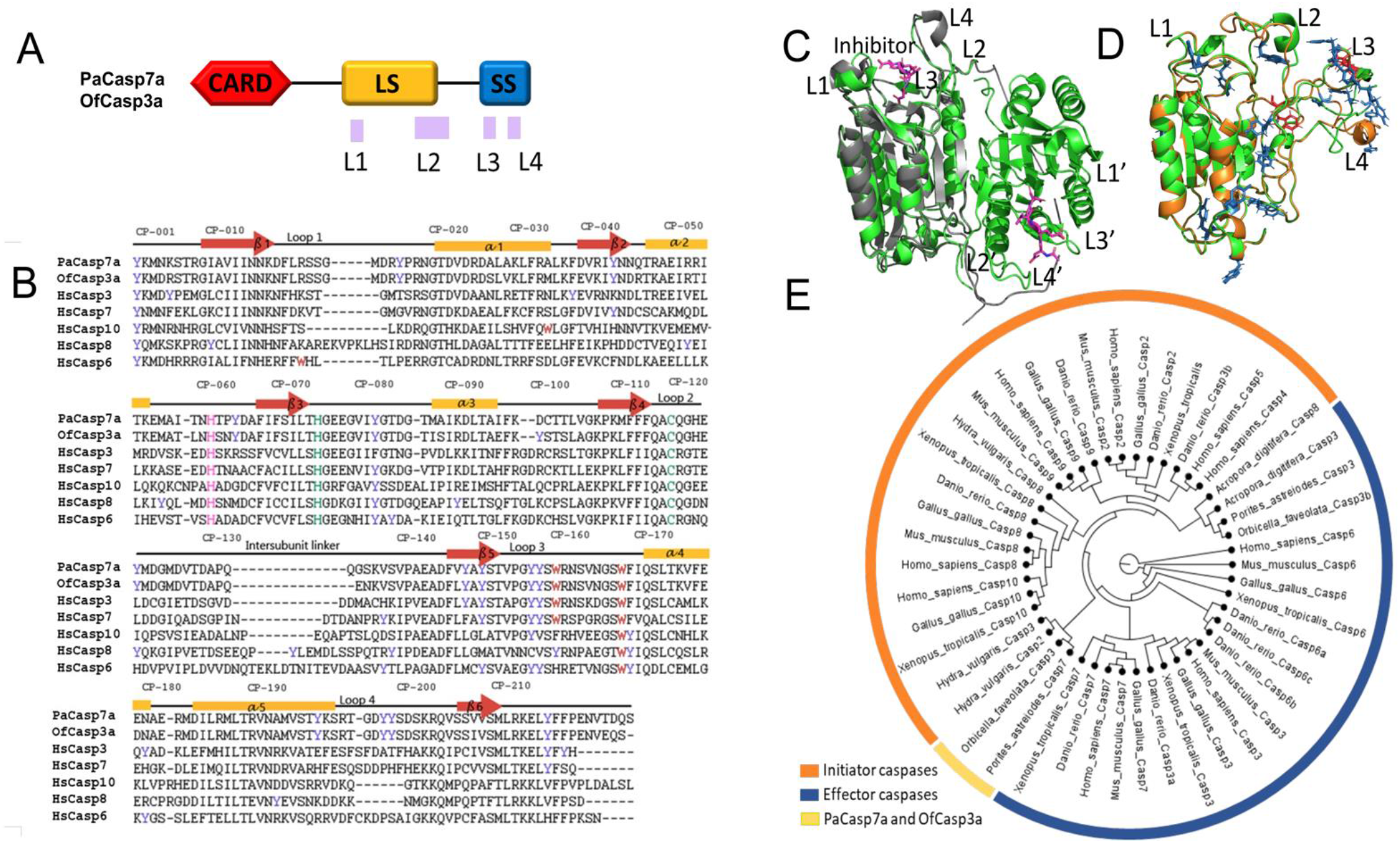
Phylogenetic relationship and caspase structure. (A) Domain organization of the protomer and location of active site loops, L1-L4, on coral caspases. The CARD domain precedes the large and the small subunit of the protease domain. (B) Multiple sequence alignment of coral caspases with human caspases showing secondary structural elements (loops, beta sheets, alpha helices) along with the common position (CP) numbers among caspases. Tyrosine residues (blue), tryptophan residues (red), active site catalytic residues His and Cys (green) and a conserved His residue (magenta) are highlighted. Sequences of the pro-domains are not shown. (C) Comparison of active dimeric PaCasp7a (green) (PDB ID: 6WI4) superimposed with HsCasp-3 protomer (gray) (PDB ID: 2J30) indicating active site loops L1-L4 on protomer 1 and L1’-L4’ on protomer 2. (D) Predicted structures of the protomer of PaCasp7a (green) and of OfCasp3a (orange) built using NMR-model structure of HsproCasp8 (PDB ID: 2k7z) with an intact intersubunit linker, as described in the text. Tryptophan (red) and tyrosine (blue) residues are highlighted. (E) Phylogenetic tree of apoptotic caspases.

All caspases have a conserved caspase-hemoglobinase (CH) fold characterized as a six-stranded β-sheet core with at least five α-helices on the protein surface (Fig.1B,C).^10^ The protomer of effector caspases is cleaved at the intersubunit linker to separate the large and small subunits, resulting in the rearrangement of several active site loops and stabilization of the active conformation. In contrast, initiator caspases are activated through dimerization, and subsequent chain cleavage is thought to stabilize the dimer. ^11,12^ Thus, the CH fold provides a basis for the evolution of regulation through oligomerization; enzyme specificity, resulting in overlapping but non-identical cellular substrates; and allosteric regulation, through both common and unique allosteric sites. As a result, caspases provide an excellent model for studying protein evolution. ^4,13^

Previous studies in the model organisms *C. elegans* and *D. melanogaster* have provided critical insights into the molecular basis of the caspase cascade; however, both organisms utilize fewer caspases than in humans, resulting in apoptosis cascades that do not reflect the characteristics of the ancestral metazoans.^14–17^ In contrast, genomic studies of cnidaria, the sister group to bilaterians comprising corals, jellyfish, and sea anemone, revealed multiple genes that were previously thought to be confined to vertebrates, thereby demonstrating the extensive gene loss in *C. elegans* and *D. melanogaster*. Indeed, the apoptotic genes in the cnidarians appear to complement the apoptotic signaling cascade observed in vertebrates, although much less is known about the cnidarian caspases compared to their counterparts in vertebrates.^18–22^ We showed recently that a caspase-7 from *Porites astreoides* (PaCasp7a), a disease-resistant reef-building coral, and a caspase-3 from *Orbicella faveolata* (OfCasp3a), a disease-sensitive reef-building coral, utilize the conserved CH fold (Fig. 1C). The presence of an N-terminal CARD on a caspase that exhibits caspase-3/-6-like enzyme specificity suggests linkage of effector caspases to death platforms in coral. ^23^ The collective data from vertebrates and invertebrates show that caspases utilize a protein fold that has been conserved for at least one billion years and that evolutionary modifications to the conformational landscape resulted in stable monomeric or dimeric subfamilies, changes in enzyme specificity, and changes in allosteric regulation. ^4,13,24–26^

The folding landscape of human caspase-3 has been studied extensively.^27,28^ Recent studies of other effector caspases from human (caspases-6 and -7) ^4^ and from zebrafish (caspases-3a and -3b) ^13^, as well as the common ancestor of effector caspases, ^4,26^ show a conserved folding landscape in which the native dimer (N_2_) is in equilibrium with at least two partially folded intermediates - I_2_ (partially folded dimer) and I (partially folded monomer) - prior to unfolding (U). The conserved landscape provides flexibility through mutations that either stabilize or destabilize the partially folded intermediates, allowing species-specific adjustments to the relative populations of partially folded states. In addition, all dimeric caspases examined to date undergo a pH-dependent conformational change, with pKa∼6, resulting in an enzymatically inactive conformation. ^4,13,29^ The reversible conformational change may provide a mechanism to regulate caspase activity through localized changes in pH.

In contrast to the dimeric caspases, very little is known about the folding landscape of monomeric caspases, and the properties have been inferred from the monomeric folding intermediate (I) of dimeric caspases. ^27^ To further understand the evolution of the monomeric subfamily of caspases, we examined the folding landscape of coral caspases from *O. faveolata* and *P. astreoides*, which are about 300 million years apart on an evolutionary timescale and about 600 million years apart from human caspases. The two proteins (PaCasp7a and OfCasp3a) are monomeric (Fig. 1D) ^23^, are orthologs of the initiator family of caspases, and share a common ancestor with effector caspases (caspases-3, -6, -7) and with initiator caspases (caspases-2, -9, -8, -10) (Fig. 1E). PaCasp7a and OfCasp3a have about 77% amino acid sequence identity with each other and about 35% sequence identity with human caspases -3/-7. ^23,26^

Here, we examined the urea-induced equilibrium unfolding of the zymogens (procaspases) of PaCasp7a and of OfCasp3a over a broad pH range, from pH 3 to pH 10.5. The results are compared to the partially folded monomeric intermediate observed during procaspase-3 (PCP-3) folding and assembly. The data show that the pH-dependent conformational change observed in the dimers is also conserved in the monomers, suggesting an evolutionarily conserved regulatory mechanism. In addition, the monomeric caspases unfold via similar mechanisms where at least one partially folded intermediate is in equilibrium with the native conformation and the unfolded state. Overall, the conformational free energy is somewhat higher than that determined for the partially folded monomeric intermediate of dimeric caspases, and the native conformation is destabilized compared to the partially folded intermediates at low and high pH. Finally, the data show that the small subunit in the protomer unfolds prior to the large subunit. Together, the data indicate that evolution of the dimeric family of caspases was important for stabilizing the small subunit within the protomer.

## Results

The monomeric procaspase PaCasp7a and OfCasp3a constructs are without an N-terminal CARD domain and consist of 260 and 258 amino acids, respectively, with a molecular weight of about ∼31 kDa (Fig 1B). Since caspase activation can be autocatalytic under certain circumstances, such as when protein concentrations are high in heterologous expression systems, we replaced the active site cysteine with a serine residue (CP-C120S) for our equilibrium unfolding studies to prevent cleavage during expression in *E. coli*. We note that we utilize the common position (CP) nomenclature, described previously, that allows amino acid position descriptions and comparisons of all caspases from any organism. ^30^ As shown in the sequence alignment in Figure 1B, the proteins begin with a uniformly conserved tyrosine (CP-001) at the N-terminus of the protease domain (that is, large subunit-intersubunit linker-small subunit). PaCasp7a and OfCasp3a have two tryptophan residues each (CP-W160 and CP-W168), and both reside in the active site loop 3 (L3), which is in the small subunit of these caspases (Fig. 1A,D). In addition, PaCasp7a and OfCasp3a have 14 and 15 tyrosine residues, respectively, well distributed in the primary sequence (Fig. 1B, blue). Finally, residues in β-strand 6, which forms the dimer interface (Fig. 1B, CP-208 to CP-213, SSV/ISML), are more similar to those of the dimeric effector caspases (CIVSML) than those of the monomeric initiator caspases (QPTFTL), suggesting that other factors also contribute to stable dimer formation.

### Urea-induced unfolding of PaCasp7a and of OfCasp3a

We examined changes in the tertiary structure at increasing urea concentrations by exciting the samples at 280 nm or at 295 nm to monitor changes in fluorescence emission of tyrosine and tryptophan residues (280 nm) or tryptophan residues only (295 nm) as a function of urea concentration. As shown in Supplemental Figure S1, native PaCasp7a (Supplemental Fig. S1, panels A and B) and OfCasp3a (Supplemental Fig. S1, panels D and E) have an emission maximum at 334 nm when excited at 280 nm and 338 nm when excited at 295 nm, similar to those of human PCP-3 (procaspase-3) and PCP-7 (procaspase-7). The fluorescence emission maxima are red-shifted to ∼347 nm in phosphate buffer containing 9 M urea, showing that the aromatic residues are exposed to solvent and that the proteins are fully unfolded under these solution conditions. Similarly, PaCasp7a (Supplemental Fig. S1, panel C) and OfCasp3a (Supplemental Fig. S1, panel F) have well-formed secondary structure that is disrupted in a 9 M urea-containing phosphate buffer, as shown by the significant change in the signal due to the loss of secondary structure, determined with far-UV circular dichroism (CD).

We examined the equilibrium unfolding of monomeric coral caspases as a function of urea concentration and over the pH range of 3 to 10.5. Since both OfCasp3a and PaCasp7a are monomers, their equilibrium unfolding is not dependent on the protein concentration, as confirmed by studies using a protein concentration range of 0.5-4 µM (data not shown). Further, refolding experiments for both proteins demonstrated reversible folding transitions. Representative data for the equilibrium unfolding of PaCasp7a and OfCasp3a at pH 3, pH 7, and pH 10 are shown in Figure 2. The data for pH 7 are described here, while those for lower and higher pHs are described below. In the case of PaCasp7a at pH 7 (Fig. 2A), one observes a pre-transition between 0 M and ∼1.5 M urea in both the fluorescence emission and CD data, showing little to no change in the signal of the native protein. The pre-transition is followed by a cooperative change in the signal between ∼1.5 M and ∼2.5 M urea, where the signal for fluorescence emission increases. Interestingly, the transition is accompanied by a near complete loss in secondary structure as shown by the decrease in CD signal. Following the cooperative change in signal, there is an apparent plateau between ∼2.5 M and ∼4.5 M urea when samples are excited at 295 nm, although one observes a modest decrease in the fluorescence signal in this region when samples are excited at 280 nm. At higher urea concentrations, one observes a second cooperative transition between ∼4.5 M and ∼6 M urea, demonstrating that the protein is unfolded at urea concentration >6 M. In comparison, OfCasp3a demonstrates a pre-transition between 0 M and ∼0.5 M urea in both the fluorescence emission and CD data, followed by a cooperative change in the signal between ∼0.5 M and ∼1.5 M urea. Like PaCasp7a, the fluorescence emission signal increases, whereas the relative signal for CD decreases during this transition. Between ∼2 M and ∼4.5 M urea, one observes a decrease in the fluorescence signal followed by a second cooperative transition between ∼5 M and ∼6 M urea, beyond which the protein is unfolded. We note that the relative fluorescence emission signal is higher when samples are excited at 295 nm compared to those excited at 280 nm and that the relative fluorescence signal is higher overall than that observed by CD for both proteins.

**Figure 2.**
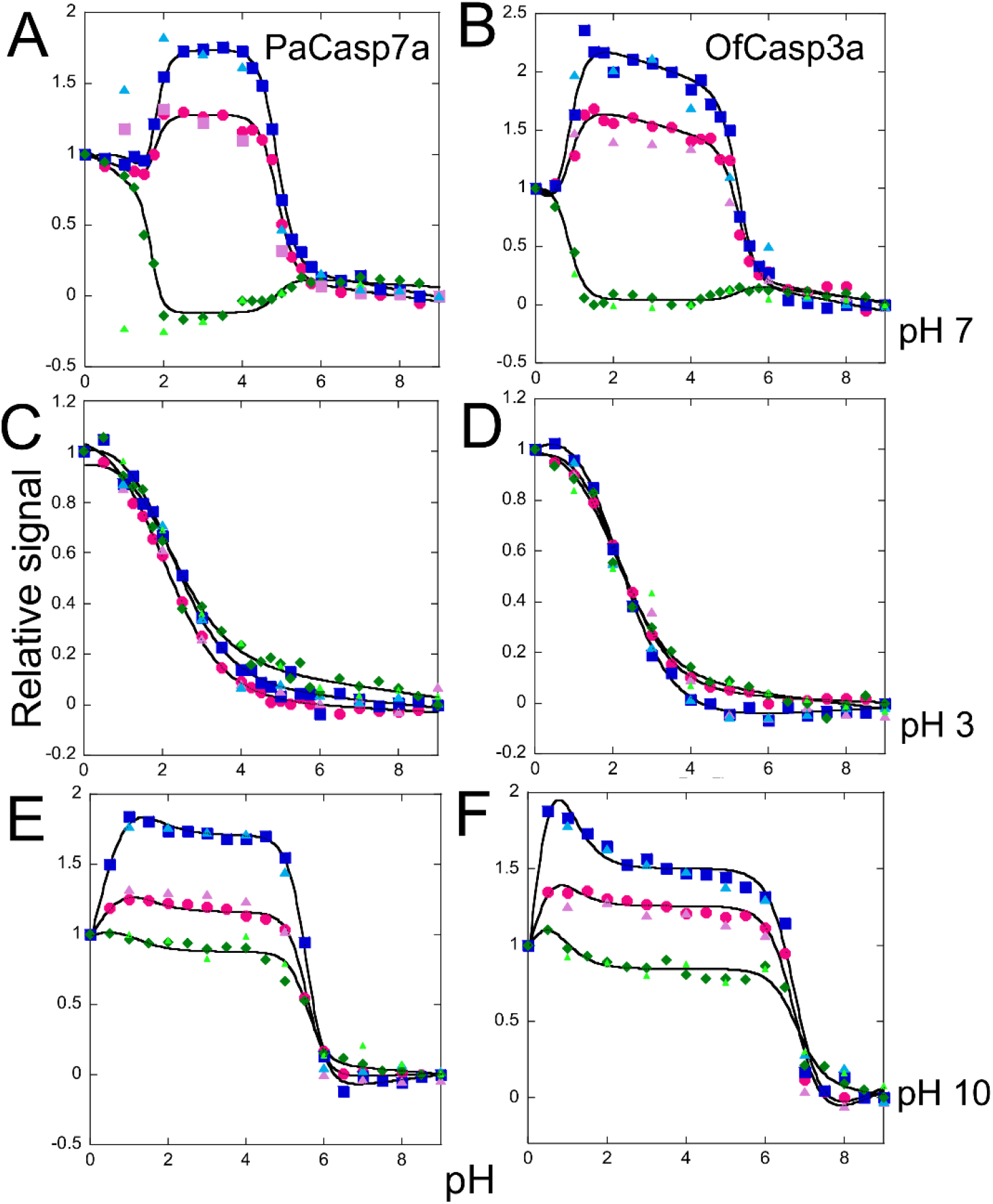
Normalized equilibrium unfolding data for PaCasp7a (left panels) at pH 7 (A), pH 3 (C) and pH 10 (E). Normalized equilibrium unfolding data for OfCasp3a (right panels) at pH 7 (B), pH 3 (D) and pH 10 (F). Colored solid symbols represent averaged raw data and solid lines through the data represent the global fits. Excitation at 280 nm-unfolding 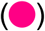 and refolding 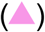, excitation at 295 nm - unfolding 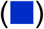 and refolding 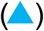, CD - unfolding 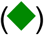 and refolding 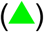

### Global fits to the equilibrium unfolding data

As described previously for the dimeric caspases^4,13,27^, the data shown in Figure 2 were fit globally to an equilibrium folding model to determine the conformational free energies. The solid lines through the data points represent the global fits. The data at pH 7 (Fig. 2 A,B) suggest that the proteins unfold via a four-state mechanism in which the native monomer (N) isomerizes to a partially folded intermediate (I_1_), followed by further unfolding through a second partially folded intermediate (I_2_) prior to unfolding (U) (N ↔ I_1_↔ I_2_↔ U) (equation 1). In general, the two partially folded intermediate conformations, I_1_ and I_2_, are observed in the data through the plateau between ∼2 M and ∼4.5 M urea. While both intermediates exhibit a higher relative fluorescence emission than the native conformation, the second intermediate, I_2_, exhibits a lower fluorescence emission than does the first intermediate, I_1_. At pH 7, for PaCasp7a, the transition from native (N) to intermediate 1 (I_1_) occurs with a free energy of 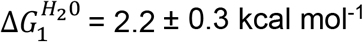, and the transition from intermediate_1_ (I_1_) to intermediate_2_ (I_2_) has a free energy of 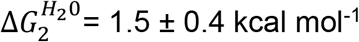. The free energy change, 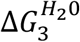, for the unfolding of the second intermediate to the unfolded state, I_2_ to U, is 11.2 ± 0.1 kcal mol^-1^, with a total conformational free energy of ∆G°_conf_ =14.9 kcal mol^-1^. The cooperativity indices m_1_, m_2,_ and m_3_ for the transitions are 1.05 ± 0.3 kcal mol^-1^M^-1^, 1.10 ± 0.4 kcal mol^-1^M^-1^, and 2.30 ± 0.4 kcal mol^-1^M^-1,^ respectively (Table I). For OfCasp3a at pH 7, the free energy change 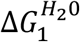 for N to I_1_ is 2.0 ± 0.4 kcal mol^-1,^ and that of I_1_ to I_2_, 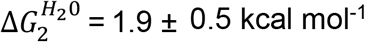. The free energy change 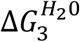, for I_2_ to U, is 13.4 ± 0.6 kcal mol^-1^ with a total conformational free energy of ∆G°_conf_ = 17.4 kcal mol^-1^. The cooperativity indices m_1_, m_2,_ and m_3_ for the transitions are 1.80 ± 0.3 kcal mol^-1^M^-1^, 1.20 ± 0.5 kcal mol^-1^M^-1^, and 2.60± 0.8 kcal mol^-1^M^-1,^ respectively (Table II). Thus, the two proteins exhibit similar conformational free energies for unfolding. Interestingly, the m-values indicate that approximately half of the buried surface area is exposed in the first two unfolding transitions. Taken together, the data suggest that the second intermediate, I_2_, is characterized by a lack of secondary structure but with substantial buried hydrophobic surface. We previously showed that the partially folded monomer of human PCP-3 has a conformational free energy change of 7.2 ± 0.5 kcal mol^-1^ at pH 7, 25°C, so both monomeric coral caspases exhibit substantially higher conformational free energy.

**Table I.** Free energy changes and co-operativity index of PaCasp7a for each unfolding transition.

| pH | Equilibrium mechanism | Free energy change<br>($\Delta G$ kcal mol<sup>-1</sup>) | m-value<br>(kcal mol<sup>-1</sup> M<sup>-1</sup>) | $\Delta G^0_{\text{total}}$<br>(kcal mol<sup>-1</sup>) | $m_{\text{total}}$<br>(kcal mol<sup>-1</sup> M<sup>-1</sup>) |
| --- | --- | --- | --- | --- | --- |
| pH 3 | I<sub>2</sub> to U | 1.6 ± 0.2 | 0.80 ± 0.1 | 1.6 ± 0.2 | 0.80 ± 0.1 |
| pH 7 | N to I<sub>1</sub> | 2.2 ± 0.3 | 1.05 ± 0.3 | 14.9 ± 0.3 | 4.45 ± 0.4 |
|  | I<sub>1</sub> to I<sub>2</sub> | 1.5 ± 0.4 | 1.10 ± 0.4 |  |  |
|  | I<sub>2</sub> to U | 11.5 ± 0.1 | 2.30 ± 0.3 |  |  |
| pH 10 | I<sub>1</sub> to I<sub>2</sub> | 0.9 ± 0.3 | 1.41 ± 0.3 | 11.9 ± 0.8 | 3.38 ± 0.3 |
|  | I<sub>2</sub> to U | 11.0 ± 0.1 | 1.97 ± 0.2 |  |  |

**Table II.** Free energy changes and co-operativity index of OfCasp3a for each unfolding transition.

| pH | Equilibrium mechanism | Free energy change<br>($\Delta G$ kcal mol<sup>-1</sup>) | m-value<br>(kcal mol<sup>-1</sup> M<sup>-1</sup>) | $\Delta G^0_{\text{total}}$<br>(kcal mol<sup>-1</sup>) | $m_{\text{total}}$<br>(kcal mol<sup>-1</sup> M<sup>-1</sup>) |
| --- | --- | --- | --- | --- | --- |
| pH 3 | I<sub>2</sub> to U | $1.7 \pm 0.3$ | $0.86 \pm 0.1$ | $1.7 \pm 0.3$ | $0.86 \pm 0.1$ |
| pH 7 | N to I<sub>1</sub> | $2.0 \pm 0.4$ | $1.80 \pm 0.3$ | $17.4 \pm 0.5$ | $5.6 \pm 0.5$ |
| | I<sub>1</sub> to I<sub>2</sub> | $1.9 \pm 0.5$ | $1.20 \pm 0.5$ | | |
| | I<sub>2</sub> to U | $13.4 \pm 0.6$ | $2.60 \pm 0.8$ | | |
| pH 10 | I<sub>1</sub> to I<sub>2</sub> | $0.8 \pm 0.2$ | $1.50 \pm 0.3$ | $12.3 \pm 0.1$ | $3.2 \pm 0.2$ |
| | I<sub>2</sub> to U | $11.5 \pm 0.1$ | $1.70 \pm 0.2$ | | |

### pH effects on equilibrium unfolding of PaCasp7a and OfCasp3a

We showed previously that pH changes are an excellent perturbant for examining the conformational landscape of caspases as both the conformational stability and oligomeric state may be affected. ^28^ In addition, the folding transitions are reversible with variations in pH but not with variations in temperature (data not shown). As shown in Figure 2 and Supplemental Figures S2-S5, we examined equilibrium folding of PaCasp7a and of OfCasp3a over the pH range of 3 to 10.5. While the data for the two extremes (pH 3 and pH 10.5) are summarized in Figure 2, the full profile is shown in Supplemental Figures S2-S5. We note that the equilibrium folding of OfCasp3a was not reversible between pH 4.5 and 6 due to protein aggregation, so data for those pHs are not considered in subsequent analyses. Similar to the folding model described at pH 7 above, the equilibrium unfolding of PaCasp7a and of OfCasp3a is well-described by a four-state model (equation 1) between pH 6 and 8. Fits of the data to the models described below are shown as the solid lines in the figures, and the ∆G°_conf_ and m-values are shown in Tables I and II and in Supplementary Tables SI and SII. Finally, the pH dependence of the conformational stability and cooperativity indices (m-value) over the entire pH range (pH 3 to pH 10.5) for both PaCasp7a and OfCasp3a are summarized in Figure 3.

**Figure 3.**
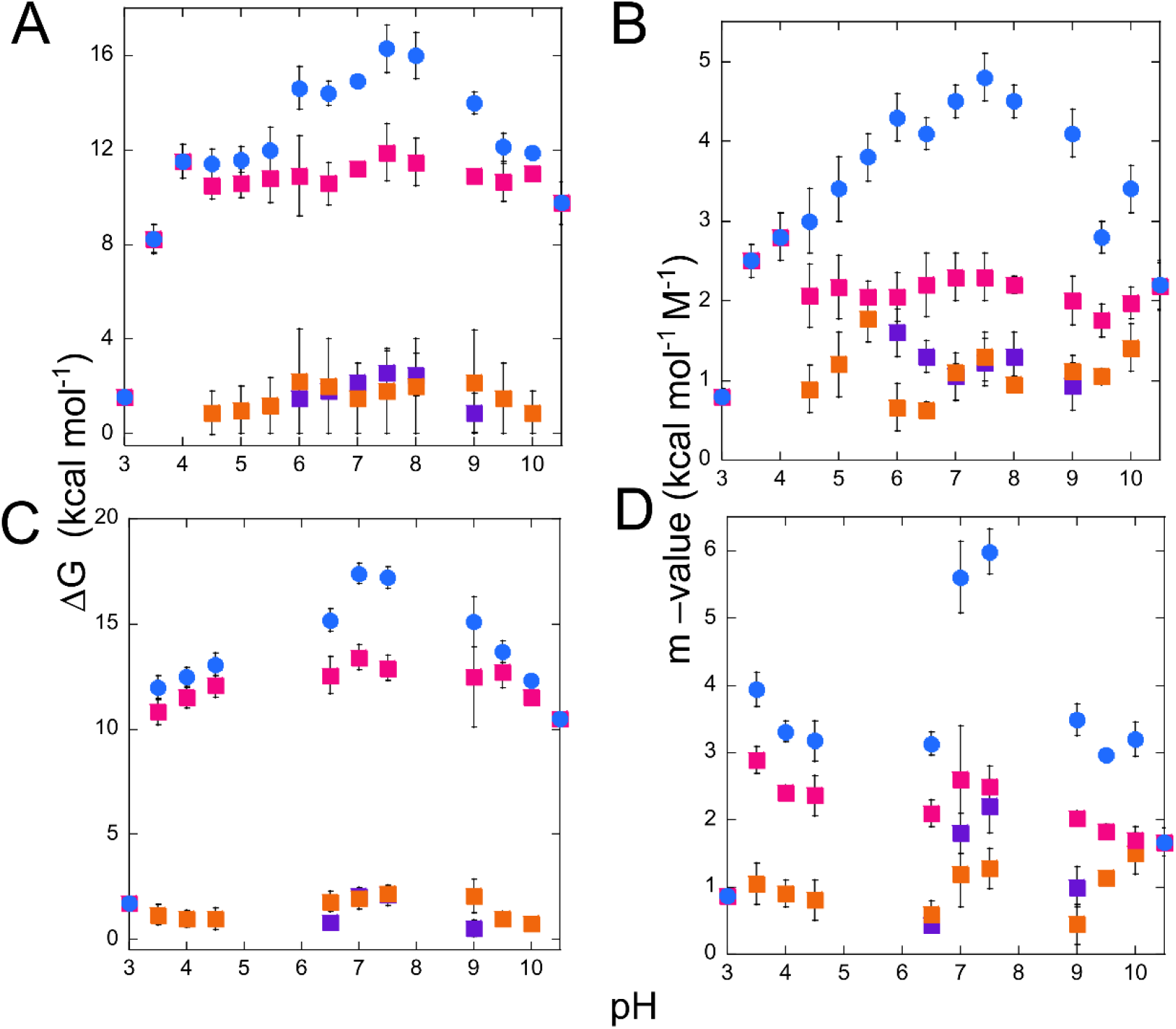
Conformational free energy as a function of pH. PaCasp7a (A,B), OfCasp3a (C,D).For panels A-D. the following symbols were used: ΔG_total_ 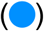, ΔG_1_ 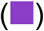, ΔG_2_ 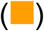, ΔG_3_ 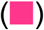. Cooperativity indices: m_total_ 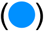, m_1_ 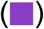, m_2_ 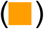, m_3_ 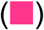. Error bars are from supplementary tables SI and SII.

First, one observes that the midpoint of the first transition shifts to lower urea concentrations as the pH is lowered, such that the native protein is destabilized below pH 6 (Supplemental Figures S2 and S4) resulting in the loss of the N-to-I_1_ transition below pH∼4.5. In other words, the “native” protein at pH 4.5 appears to comprise the I_1_ partially folded state, so the data at pH 4.5 were best-fit to a three-state folding model (I_1_↔I_2_↔U, equation 2). As the pH is lowered further, the mid-point of the transition of I_1_ to I_2_ decreases such that the transition of I_1_ to I_2_ is lost, and the data are best-fit to a two-state equilibrium folding model at pH 3 (I_2_↔U, equation 3) (Fig. 2C,D and Supplemental Figures S2 and S4). A similar process is observed at higher pH. As the pH is increased above pH 9, the midpoint of the N to I_1_ transition shifts to lower urea concentrations such that N is destabilized at pH>9, and the data are best fit first to a three-state equilibrium unfolding model (I_1_↔I_2_↔U, equation 2, pH 9.5-10), then to a two-state equilibrium folding model at pH 10.5 (I_2_↔U, equation 3) (Figure 2D,E and Supplemental Figures S2 and S4). For both PaCasp7a and OfCasp3a, the total conformational free energies demonstrate that the proteins are most stable near physiological pH and that the stability decreases at both lower and higher pH due to destabilizing first the native conformation then the first folding intermediate, I_1_.

Overall, the data show that the higher stability near physiological pH, between pH 6 and ∼8, with a ∆G°_conf_∼16 kcal mol^-1^, is largely due to the contribution of the partially folded intermediate, I_2_, with 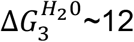 kcal mol^-1^. While the stability of the native conformation and of the partially folded intermediate, I_1_, decrease at lower and higher pH, the stability of I_2_ appears to be largely independent of pH and decreases only below pH 4 (Fig. 3A,C). The cooperativity index (m-value) is correlated to the change in accessible surface area (ΔASA) exposed to solvent during each unfolding transition. The total m-value (m_total_) is similar for PaCasp7a and OfCasp3a, indicating a similar exposure of hydrophobic area during each unfolding transition (Fig. 3B,D). Together, the data suggest that partially folded conformations are well-populated at lower and higher pHs and that the two monomeric caspases exist in their native conformation only between the pH range of 6 to 8.

The changes in the total conformational free energy shown in Figure 3 for both proteins are compared in Figure 4A and show three transitions over the pH range of 3 to 10.5, mostly due to the changes in the populations of N and I_1_ at lower and higher pH as well as the lower stability of I_2_ below pH 4. In order to further examine the conformational changes due to pH, we performed titrations of native protein versus pH, in the absence of urea, and measured changes in secondary structure (Fig. 4B). We also measured changes in average emission wavelength for both proteins following excitation at 280 nm or at 295 nm (Fig. 4C,D). As described above for urea-induced equilibrium unfolding, the proteins exhibit a substantial loss of secondary structure during the transition of N to I_1_. Likewise, one observes a substantial loss of secondary structure when the native protein is titrated from pH 7 to pH 6 (Fig. 4B). While the secondary structure decreases at higher pH, the signal loss is lower than that observed at lower pH. In terms of the tertiary structure, the average emission wavelength is relatively constant at pH>6, and one observes a cooperative decrease below pH 6. We fit the data in Figure 4 to determine the pKa values for each transition, as described in Methods, and the fits are shown as the dashed lines in the figure. Although the data for OfCasp3a were not well-determined due to the lack of data at all pH values, we note that both proteins exhibit similar transitions and have identical pKa values. The fits to the total free energy, ∆G°_conf_ (Fig 4A) can be characterized by three pKa values between pH 3 and pH 10.5 for PaCasp7a and for OfCasp3a. We observe that, on decreasing the pH from 7 to 5, there is a decrease in the overall stability characterized by an estimated pKa of ∼5.7. The transition correlates with the transition observed in the average emission wavelength of the native PaCasp7a and OfCasp3a (Fig 4C,D), which can be characterized by an estimated pKa ∼5.3. Changes in the secondary structure for PaCasp7a and OfCasp3a show a sharp transition (Fig. 4B), with an estimated pKa of 6.1. We previously demonstrated that the dimers of PCP-3, PCP-6, and DrPCP-3b undergo a pH-dependent conformational change, with a pKa of 5.7. ^4,13,29^ Together, the data suggest that the decrease in the average emission wavelength and the conformational stability report the same pH-dependent conformational change in the monomeric coral caspases. That is, titration of one or more residues affects the transition of the native conformation to a partially folded intermediate. In the dimeric caspases, the transition is accompanied by a loss of enzymatic activity, although the protein remains dimeric. In the monomeric caspases, shown here, the transition is accompanied by a loss in secondary structure and a blue-shift in fluorescence emission. In addition, the proteins undergo a further transition with pKa∼3.3. The data show that the transition is due to titration of the partially folded intermediate, I_2_. Interestingly, the transition that occurs at higher pH, with pKa∼9.3, is not accompanied by a blue-shift in fluorescence emission even though the secondary structure is destabilized. The simplest model suggests that the native conformation is destabilized relative to the partially folded intermediates at both lower and higher pH. If this is true, then the fluorescence emission at higher pH may not reflect the blue-shift until all secondary structure is lost, indicating a broader pH range would be required at higher pHs.

**Figure 4.**
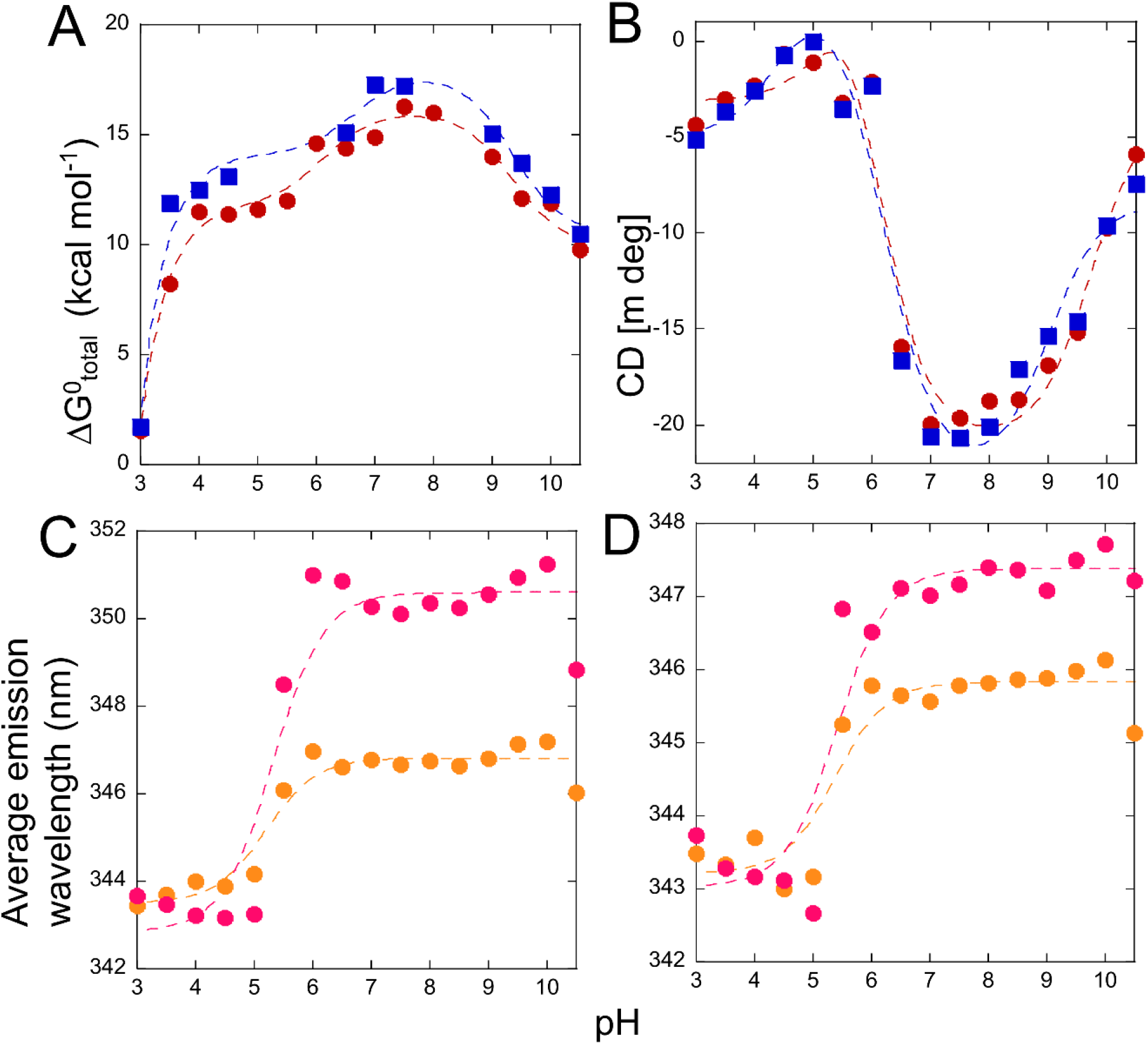
Effect of pH on monomeric coral caspases. (A) Comparison of the ∆G^0^_total_ for PaCasp7a 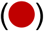 and OfCasp3a 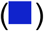. (B) Change in CD signal for native PaCasp7a 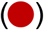 and native OfCasp3a 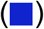. (C) Change in average emission wavelength versus pH for PaCasp7a (C) and for OfCasp3a (D). For panels C and D, samples were excited at 280nm 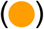 or 295nm 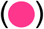. For A-D, dashed lines represent fits to the data to determine the pKa of the transitions, as described in the text.

### Molecular dynamics (MD) simulations and limited proteolysis show the small subunit is unstable

The equilibrium unfolding data described above demonstrate that the proteins have substantial conformational free energy between pH 5 and 10, with ∆G°_conf ∼_12-17 kcal mol^-1^ (Fig. 4A). Yet, based on data from urea-induced unfolding (Fig. 2) and from far-UV CD spectra (Fig. 4B), the proteins lose most of their secondary structure below pH 7 and above pH ∼10. In addition, the proteins exhibit a cooperative unfolding transition following the apparent loss of secondary structure (Fig. 2), with a corresponding cooperativity index (m-value) that suggests a substantial hydrophobic core that remains solvent inaccessible. In contrast to the CD data, the average emission wavelength (AEW) does not change with pH>5, where the AEW remains constant at ∼347 nm, which suggests that the tryptophan residues are in a hydrophilic environment (Fig. 4C,D). To further examine how the proteins may lose secondary structure at lower and higher pHs while retaining a partially folded conformation, we performed MD simulations at several pHs and in the presence and absence of 5 M urea.

PaCasp7a and OfCasp3a protomers were modeled using the procaspase-8 structure determined by NMR (PDB ID: 2k7z),^31^ since no other structures are available for monomeric procaspases. The inter-subunit linker in PaCasp7a and OfCasp3a is intact as is observed in the model (Fig. 1B, Fig. S6). All ß-strands in the hydrophobic core of the protein are well-formed except for ß6. Indeed, ß6 forms multiple contacts in the interface of the dimeric effector caspases but is less well-packed in the monomers.

MD simulations were carried out for 200 ns, which is sufficient time to observe changes in the protein conformation in the presence of urea. Data for PaCasp7a and for OfCasp3a in the absence of urea offer a baseline against which we compared data for both proteins in the presence of 5 M urea. The unfolding of proteins during urea denaturation eventually exposes the hydrophobic core to the aqueous environment, resulting in the loss of hydrophobic contacts. The solvent-accessible surface area (SASA) can be used to determine changes in the solvent accessibility throughout the simulation ^32^. Overall, the native PaCasp7a and OfCasp3a at pH 4, 7, and 9 in the absence of urea show a similar SASA of ∼140 nm^2^. However, in the presence of 5 M urea, the SASA of PaCasp7a and of OfCasp3a is much larger, ∼195 nm^2^, due to higher fluctuations as a result of unfolding (Fig. 5A,C). To examine residue-level changes, we computed root mean square fluctuations (RMSF) for both proteins in water vs 5 M urea. The RMSF data for proteins in water were subtracted from those in 5 M urea to determine regions of the protein with increased fluctuations in urea. In addition, the experiments were performed at pH 4, pH 7, and pH 9. The data show that, in the absence of urea, PaCasp7a and OfCasp3a show minimal fluctuations overall and that the active site loops as well as the N- and C-termini show the largest fluctuations (Supplemental Fig. S6). However, in the presence of 5 M urea, one observes elevated fluctuations in the region of helices 2 and 3, the inter-subunit linker, and the small subunit, particularly helices 4 and 5 (Fig. 5C,D and Supplemental Fig. S6 B,D). We examined snapshots of the protein structures over the course of the simulations in 5 M urea, and the data show that the increased fluctuations in the small subunit as well as in helices 2 and 3 correlated with unfolding (Fig. 5B,D and Supplemental Fig. S6). At the end of the simulation, the small subunit is largely unfolded and helices 2 and 3 are pulled away from the protein, exposing the core ß-strands (Fig.S7). Similar results to those at pH 7 were observed at both low and high pH, except that the unfolding of the small subunit occurred earlier in the simulation, which may correlate with the lower ∆G°_conf_ observed in the equilibrium unfolding studies described above. Altogether, the data show that both proteins behave similarly, wherein the small subunit unfolds first while helices 2 and 3 separate from the body of the large subunit.

**Figure 5.**
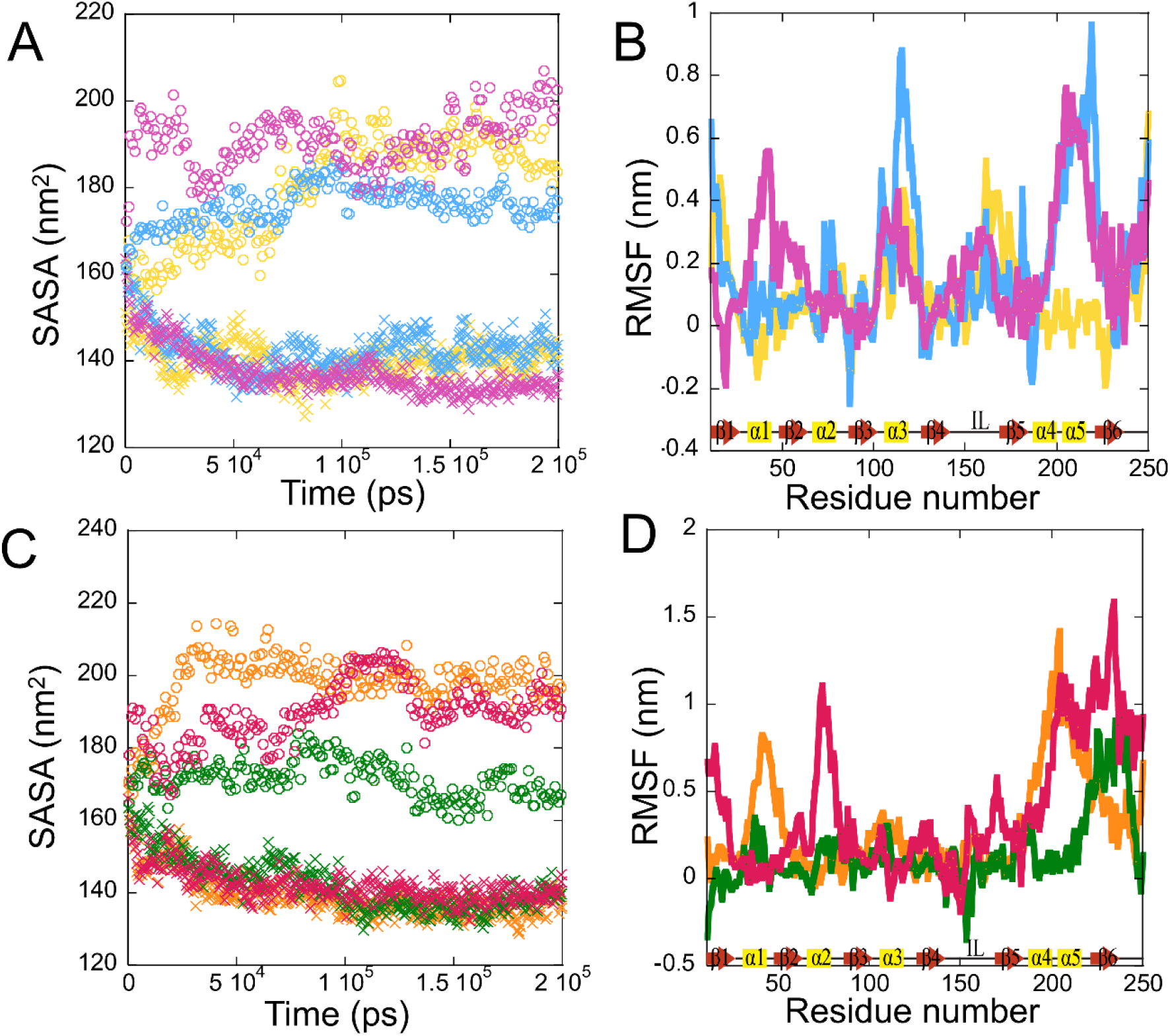
MD simulations of PaCasp7a and OfCasp3a. Panels A and C, solvent accessible surface area (SASA) of PaCasp7a (A) and of OfCasp3a (C) in water (crosses) and in 5 M urea (open circles). Panels B and D, ∆RMSF plots of 200 ns MD simulations with baseline correction as described in the text for PaCasp7a (B) and for OfCasp3a (D). For panels A and B, the following colors were used: pH 4 (yellow), pH 7 (blue), pH9 (magenta). For panels C and D, the following colors were used: pH 4 (orange), pH 7 (green), pH9 (pink).

To further examine conformational changes in PaCasp7a and in OfCasp3a due to changes in pH, we performed limited proteolysis studies at pH 7 and 9 using trypsin. The results show that trypsin cleaves the proteins in discrete regions to generate several products (Supplemental Fig S8A-D). One observes that PaCasp7a and OfCasp3a are cleaved initially to produce slightly smaller, although mostly intact proteins, suggesting cleavages near the termini (Band 2 in Supplemental Fig. S8A,C). The proteins are then further cleaved to generate fragments of 26 (Band 2), 20 (Band 3), 18 (Band 4), 15 (Band 5), and 9 kDa (Band 6). We analyzed the digests by MALDI-TOF mass spectrometry, and the results show that the first two cleavages occur at CP-R161 and GP9-R02 (see Fig. 1A), respectively, generating the 26 kDa and 20 kDa fragments. CP-R161 is located in active site loop 3 (Fig. 6). In the protomer, loop 3 is disordered, so CP-R161 is solvent exposed (Fig. 6A). In contrast, loop 3 forms the substrate binding pocket in the active dimer, so CP-R161 is more buried (Fig. 6B). In addition, GP9-R02 is located between α-helices 4 and 5, again suggesting that the two helices in the small subunit are unstable. Further cleavages of the C-terminus, inter-subunit linker, and CP-R018 on Loop 1 (R64) generate the remaining fragments. It has been reported that CP-R161 in the dimeric caspase-3 is cleaved by trypsin with a t_½_∼75 minutes at pH 7.2 and 25 °C. ^29^ However, CP-R161 is cleaved in both PaCasp7a and OfCasp3a, with a t_½_∼ 10 minutes. When considered with the result that most of the cleavages occur in the small subunit, then data suggest that the small subunit is unstable in the protomer. We note that the same cleavages occur at pH 9 as observed at pH 7, but with faster kinetics, again supporting the conclusion that the small subunit is less stable at higher pH (Supplemental Fig. S8B,D). Together, the results from MD simulations and limited proteolysis show that the small subunit of the monomeric coral caspases is unstable and unfolds prior to the large subunit.

**Figure 6.**
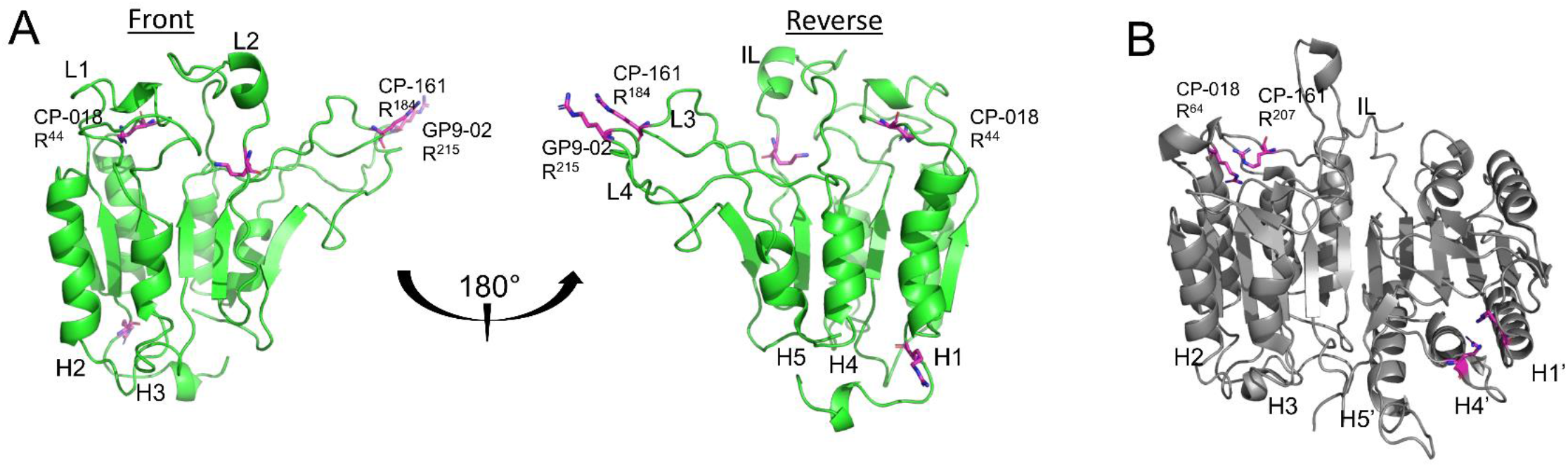
Limited trypsin proteolysis of PaCasp7a. (A) PaCasp7a protomer modeled using PDB ID:2k7z (B) Dimeric human caspase-3 (PDB ID:2J30). Common trypsin cleavage sites between the two structures are highlighted (magenta) and generate fragments as described in the text. ɑ-helices (H1-H5) (on the first protomer).

## Discussion

In this study, we characterized the equilibrium unfolding of monomeric caspases from two species of coral that are ∼300 million years distant on an evolutionary timescale. Equilibrium unfolding data suggest that PaCasp7a and OfCasp3a unfold via two partially folded intermediates in equilibrium with the native and the unfolded state, showing maximum stability of ∼17 kcal mol^-1^ near the physiological pH range (pH 6 to pH 8). The native state is destabilized outside of this pH range, and at the extreme pH the intermediate state (I_1_) is also destabilized. The conformational free energy of monomeric caspases (∼15 kcal mol^-1^) is higher than that of the partially unfolded monomeric intermediates observed during the folding and assembly of dimers, (∼7 kcal mol^-1^).

In the absence of urea, CD data indicate a substantial loss of secondary structure below pH 7 and above pH 8, yet the proteins have a conformational free energy of ∼12 kcal mol^-1^. The results suggest that intermediate I_2_ is characterized as having a strong core with buried hydrophobic contacts. The MD simulations in the presence of urea als reveal that the small subunit is unstable and unfolds prior to the large subunit, while helices 2 and 3 and expose the core β-strands. Data from limited trypsin proteolysis also indicate that the small subunit is unstable in the protomer. The lower stability of the small subunit in the protomer suggests the importance of forming the dimer since β-strand 6 forms several inter-protomer contacts in the dimer.

In general, the conformational stability decreases when the pH is reduced from 7 to 5, and the transition occurs with a pKa of ∼5.9. Similar pH-dependent conformational changes in the dimeric family of caspases have been reported^4,13,29^. Together, the data suggest that an evolutionarily conserved site is titrated with a pKa∼6, indicating that either a histidine or an acidic residue controls an important conformational switch. For example, in the dimer, the switch results in the transition of the native dimer (N_2_) to an enzymatically inactive conformation, I_2_. ^4,13,27,29^ In the monomers, the switch also destabilizes the native conformation relative to an intermediate, which may affect the formation of the dimer. Nevertheless, the conformational switch observed in all caspases could be used as a regulatory mechanism to control the activity of caspases in cells, either through changes in the active site (dimeric caspases) or through destabilizing the protomer in monomeric caspases.

The folding landscape of the dimeric caspase subfamily is conserved and provides flexibility through changes in the population distributions of two folding intermediates, a monomeric (I) and a dimeric (I_2_) intermediate. ^4,13,27,28^ Changes in the population of the two intermediates allow for species-specific modulations in overall protein stability. The caspase subfamilies evolved from a single ancestral scaffold with a conserved caspase-hemoglobinase fold that has been retained for over 650 million years, making them an excellent model for studying protein evolution. Previously, the folding mechanism of the resurrected common ancestor (CA) of the dimeric subfamily has been compared with the extant human caspases-3/-6/-7, and it was shown that the conformational free energy of the dimeric family ranges from 15 to 34 kcal/mol, ^4^ but there was little information regarding the monomer in the absence of dimerization.

PaCasp7a and OfCasp3a are evolutionarily distant from human caspases (600 million years), yet they provide a foundation for understanding the folding landscape of the monomeric family of caspases and the evolutionary events that led to the stable dimer. When considering the results described here with those of the dimeric caspases, we suggest a combined model where dimer formation results in the stabilization of the small subunit of the protomer. Evolutionary changes in the protomer that stabilize the small subunit may facilitate dimerization by decreasing fluctuations in regions of the small subunit that make up the bulk of contacts in the dimer.

## Materials and Methods

### Cloning, protein expression, and purification

Gibson cloning was used to clone PaCasp7a and OfCasp3a without their N-terminal CARD domain, and the active site cysteine was mutated to serine using site-directed mutagenesis, as described previously. ^33^ The inactive PaCasp7a and OfCasp3a zymogens were cloned into a pET-11a expression vector with a C-terminal HisTag, and all proteins were expressed in *E. coli* BL21 (DE3) pLysS cells and purified as previously described. ^27^

### Phylogenetic analysis

Caspase sequences of representative species listed in Table SIII were retrieved from CaspBase,^30^ coupled with top hits from BLAST and HMMER,^34^ and multiple sequence alignments were generated using PROMALS3D^35^ The optimal model of evolution for constructing a phylogenetic tree was determined using IQ-TREE, and the tree was constructed using the maximum likelihood approach using the Jones-Taylor Thornton model and distribution.^36^ As a test for phylogeny, the tree was bootstrapped 1000 times.

### Stock solutions and sample preparation for equilibrium unfolding

Equilibrium unfolding experiments were carried out as previously described. ^37^ Briefly, urea stock solutions (10 M) were prepared in citrate buffer (50 mM sodium citrate/citric acid, pH 3 to pH 5.5, 1 mM DTT), potassium phosphate buffer (50 mM potassium phosphate monobasic/potassium phosphate dibasic, pH 6.0-8.0, 1 mM DTT), and glycine-NaOH buffer (50 mM glycine/NaOH, pH 9 to pH 10.5, 1 mM DTT). For unfolding reactions, samples were prepared in the corresponding buffer with final urea concentrations between 0 M and 9 M. For refolding reactions, the protein was first incubated in a 9 M urea-containing buffer for ∼6 hours at 25 °C. The unfolded protein was then diluted with the corresponding buffer and urea such that the final urea concentrations were between 0.5 M and 9 M. All solutions were prepared fresh for each experiment and were filtered (0.22 μm pore size) before use. Final protein concentrations of 0.5 µM - 4 µM were used. The samples were incubated at 25°C for a minimum of 16 hours to allow for equilibration.

### Fluorescence emission and CD measurements

Fluorescence emission was measured using a PTI C-61 spectrofluorometer (Photon Technology International) from 310 to 410 nm following excitation at 280 or 295 nm. Excitation at 280 nm follows tyrosinyl and tryptophanyl fluorescence emission, whereas excitation at 295 nm follows the tryptophanyl fluorescence emission. CD data were measured using a J-1500 CD spectropolarimeter (Jasco) between 210 and 260 nm.

Fluorescence and CD spectra were measured using a 1 cm path length cuvette and constant temperature (25 °C). All data were corrected for buffer background.

### Data analysis and global fits to the equilibrium unfolding data

All data were fit globally as described previously.^27,28,37^ Briefly, equilibrium unfolding data were collected for fluorescence emission (two excitations) and far-UV CD, and at several protein concentrations, between pH 3 and 10.5 for both PaCasp7a and OfCasp3a, providing six data sets at each pH. The data were fit globally to a two-state, three-state, or a four-state equilibrium folding model, as described. ^37^ Data collected between pH 6 and pH 8 were fit to a four-state equilibrium folding model shown in equation 1. In this model, the native protein unfolds via the presence of two monomeric intermediate species, I_1_ and I_2_, prior to forming the the unfolded state. The equilibrium constants K_1_, K_2_, and K_3_ relate to equilibrium constants at respective unfolding steps.

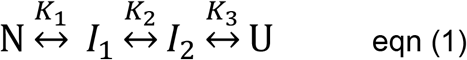

Data collected between pH 4.5 and 6, as well as between pH 9 and pH 10.5, were fit to a 3-state equilibrium folding model as described previously ^27,28,37^ and shown in Equation 2. In this model, the intermediate state (I_1_) unfolds to the intermediate (I_2_) state before complete unfolding.

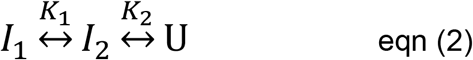

Data collected for PaCasp7a between pH 3 and pH 4 were fit to a two-state equilibrium folding model as described ^37^ and shown in equation 3,

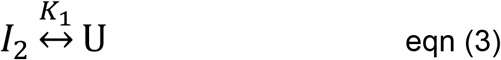

### Limited trypsin proteolysis

Proteins (6 µM) were digested with 0.5 ng/µL of trypsin in a buffer of 50 mM potassium phosphate, pH 7, 1 mM DTT at 25 °C. On addition of trypsin, aliquots were withdrawn at prescribed time intervals, and reactions were inhibited by adding SDS-PAGE buffer and heating to 100 °C for 10 minutes. The samples were frozen at −20°C until analyzed. Samples were analyzed with 4-20% SDS-PAGE gradient gels.

### MALDI-TOF mass spectrometry

Protein bands generated by limited trypsin proteolysis and resolved on SDS-PAGE gels were excised and destained with a solution of acetonitrile and 50 mM ammonium bicarbonate (1:1 v/v) for about 4 hours. The gel fragments were crushed in microcentrifuge tubes, and the proteins were extracted with a solution of formic acid/water/2-propanol (1:2:3 v/v/v) for 8 hours at room temperature. After extraction, samples were centrifuged, and the supernatant was lyophilized and redissolved in matrix solution (formic acid/water/2-propanol saturated with sinapinic acid). Protein was retrieved for MS analysis using the dried-drop method of matrix crystallization ^38^ and then analyzed using MALDI-MS (Axima Assurance Linear MALDI TOF). Laser power was optimized and 5 shots were collected, with beam blanking turned on to blank high intensity matrix peaks. Pulsed extraction was set to 2/3 maximum molecular weight. Protein standards of known masses were used to calibrate time of flight. Data were processed using data smoothing, baseline subtraction and threshold detection. Mass analysis of protein digests and cleavage sites was performed using the protein prospector software. ^39^

### MD simulations

PaCasp7a and OfCasp3a monomers were modelled using the Swiss-modeler program that uses homology modeling algorithm with user-defined templates. ^40^ PaCasp7a and OfCasp3a sequences were threaded onto the NMR structure of procaspase-8 (PDB ID:2k7z), and the modeled proteins were protonated according to the calculated ionization states of titratable groups at the specified pH using the H++ server ^41^. The force field parameters for urea were obtained as described previously, ^42^ and the urea molecule was built using the Avogadro software. ^43^ A cubic box of 6 × 6 × 6 nm^3^ containing 560 molecules of urea was generated. The system was subjected to energy minimization with the steepest-descent algorithm down to a maximum gradient of 2000 kJ mol^−1^ nm^−1^ and was simulated for 1 ns with annealing from 300 to 0K under an isotropic pressure of 100 bar. The system was then relaxed for 1 ns at standard pressure, heated from 0 to 300K, then simulated for 1 ns at 300K. The size of the box at the completion of this equilibration technique was 3×3×3 nm^3^. Using the Nosé-Hoover coupling algorithm, ^44,45^ we performed 100 ps MD simulations using the NVT (constant volume and temperature) and NPT (constant pressure and temperature) ensemble at 300K starting from the relaxed box. After heating the simulated system to 300K, a production run for each protein was conducted for 100 ps using the NPT ensemble. MD simulations were performed for 200 ns with GROMACS using the Amber99 force field and the TIP3P water model as described. ^46^

## Abbreviations

DED: death-effector domain
CARD: caspase activation and recruitment domain
DISC: death inducing signaling complex
CH: caspase-hemoglobinase
PCP: procaspase
CP: common position
SASA: solvent accessible surface area
AEW: average emission wavelength
CD: circular dichroism
MD: molecular dynamics
NMR: nuclear magnetic resonance
RMSF: root mean square fluctuation
DTT: dithiothreitol
IPTG: isopropyl ß-D-1-thiogalactopyranoside.

## Supplementary material description

Figure S1. Fluorescence emission and circular dichroism spectra of PaCasp7a and of OfCasp3a at pH 7, 25°C.

Figure S2. Normalized CD and equilibrium unfolding data for PaCasp7a from pH 3 to pH 10.5.

Figure S3. Fraction of species of PaCasp7a equilibrium unfolding as a function of urea concentration from pH 3 to pH 10.5.

Figure S4. Normalized CD and equilibrium unfolding data for OfCasp3a from pH 3 to pH 10.5.

Figure S5. Fraction of species of OfCasp3a equilibrium unfolding as a function of urea concentration from pH 3 to pH 10.5.

Figure S6. RMSF values from MD simulations mapped onto modeled structure of PaCasp7a.

Figure S7. Urea MD snapshots of PaCasp7a at pH 4, 7 and 9.

Figure S8. Limited trypsin proteolysis of PaCasp7a and OfCasp3a at pH 7 and 9.

Table SI. Summary of free energy changes and co-operativity index of PaCasp7a

Table SII. Summary of free energy changes and co-operativity index of OfCasp3a.

Table SIII. List of caspases used in the phylogenetic analysis.

## Data availability

All data are contained in the article and supporting information.

## Author contributions

I.J. and A.C.C conceptualization; I.J. and A.C.C methodology; I.J. investigation; I.J. and A.C.C writing-review and editing; A.C.C supervision.

## Funding and additional information

This work was supported by a grant from the National Institutes of Health (grant number: GM127654 [to A. C. C.]). The content is solely the responsibility of the authors and does not necessarily represent the official views of the National Institutes of Health.

