## Supplemental Data for "Sequential unfolding mechanisms of monomeric caspases"

**Running title**: pH-effects on the stability of monomeric caspases

**This file includes:**

Supplementary figures S1-S8

Supplementary tables SI-SIII

**Supplementary figure S1.**

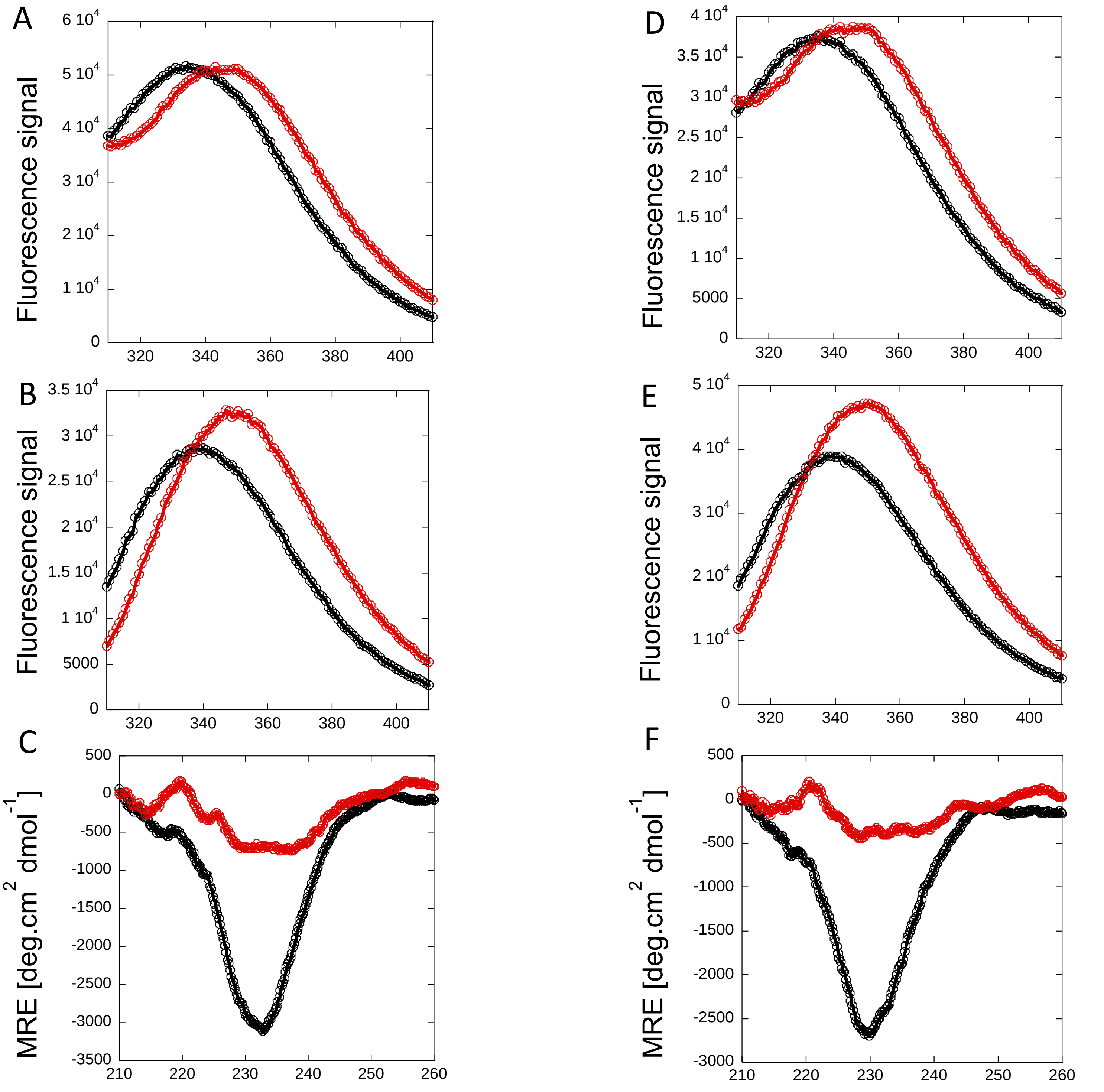


**Figure S1.** Fluorescence emission and circular dichroism spectra of PaCasp7a and of OfCasp3a at pH 7, 25°C. Fluorescence emission spectra of PaCasp7a following excitation at 280 nm (A) or 295 nm (B), and CD spectra (C). Fluorescence emission spectra of OfCasp3a following excitation at 280 nm (D) or 295 nm (E), and CD spectra (F). For A-F, the following symbols were used: proteins in buffer containing zero urea (●), or 9M urea (●)

**Supplementary figure S2 .**

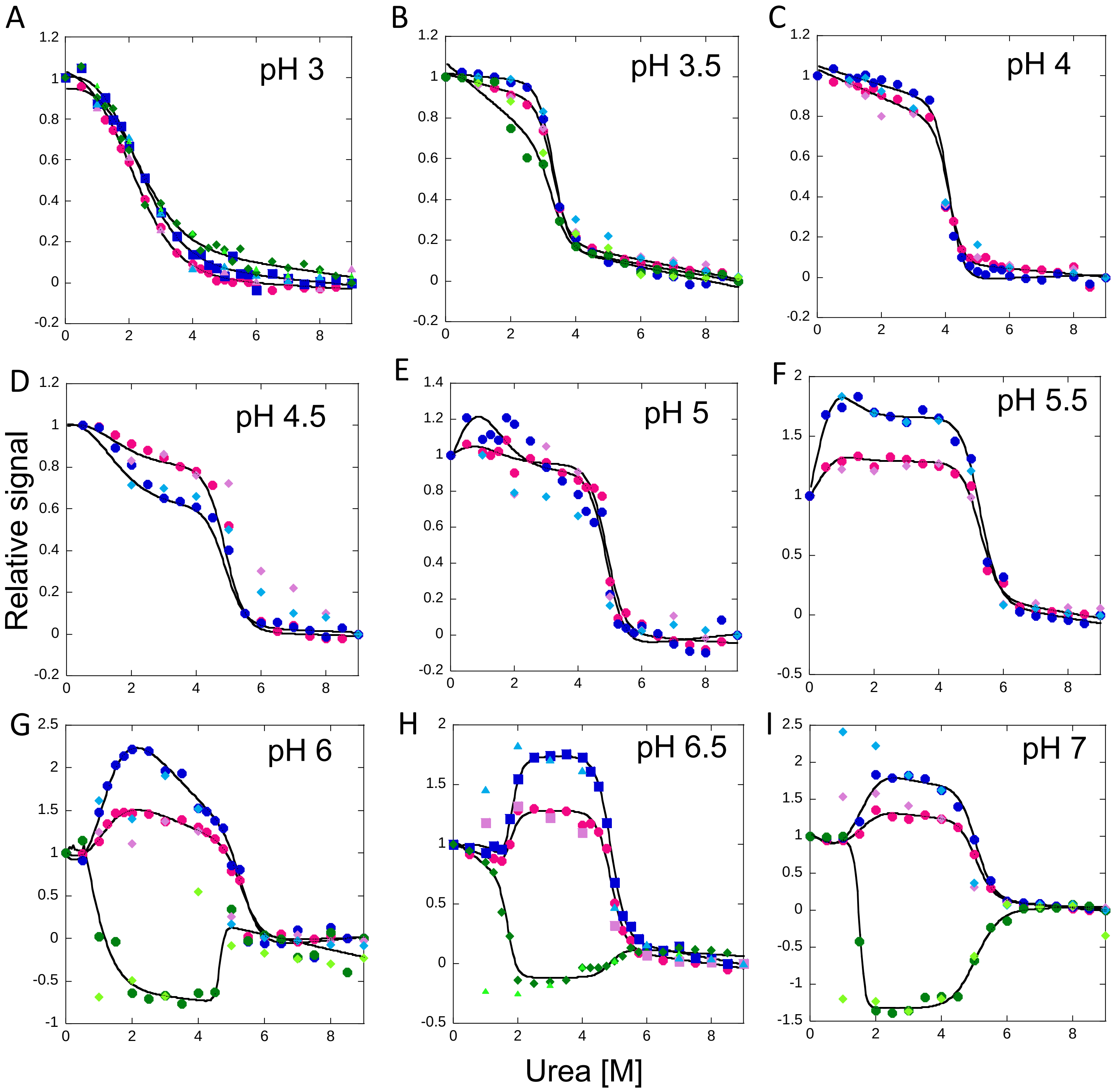

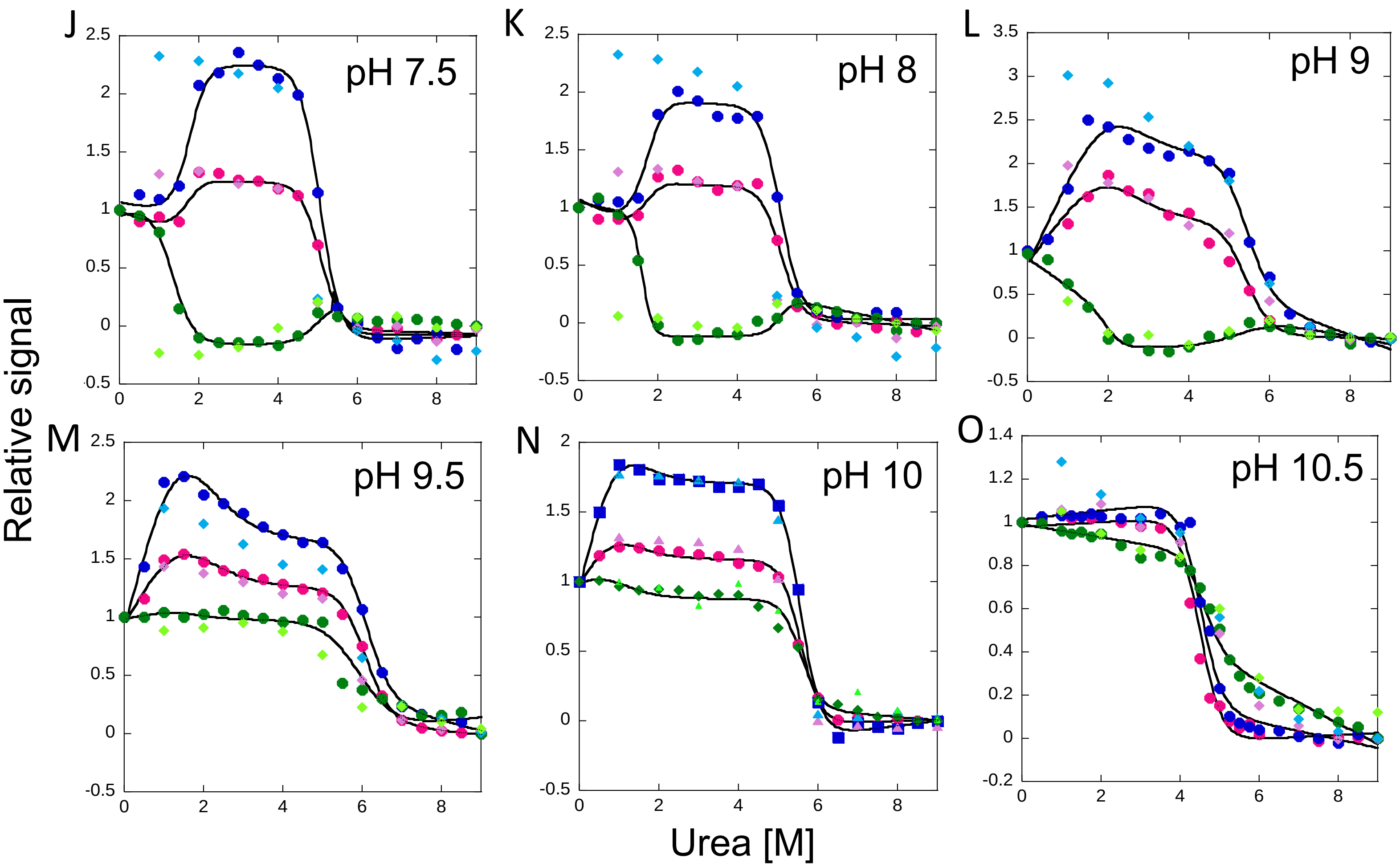


**Figure S2.** Normalized CD and equilibrium unfolding data for PaCasp7a from pH 3 to pH 10.5. Colored solid symbols represent averaged raw data and solid lines through the data represent the global fits, as described in the main text. For A-O, the following symbols were used: excitation at 280 nm- unfolding(●) and refolding(◆) , excitation at 295 nm - unfolding (●) and refolding (◆), CD - unfolding (●) and refolding (◆)

**Supplementary figure S3 .**

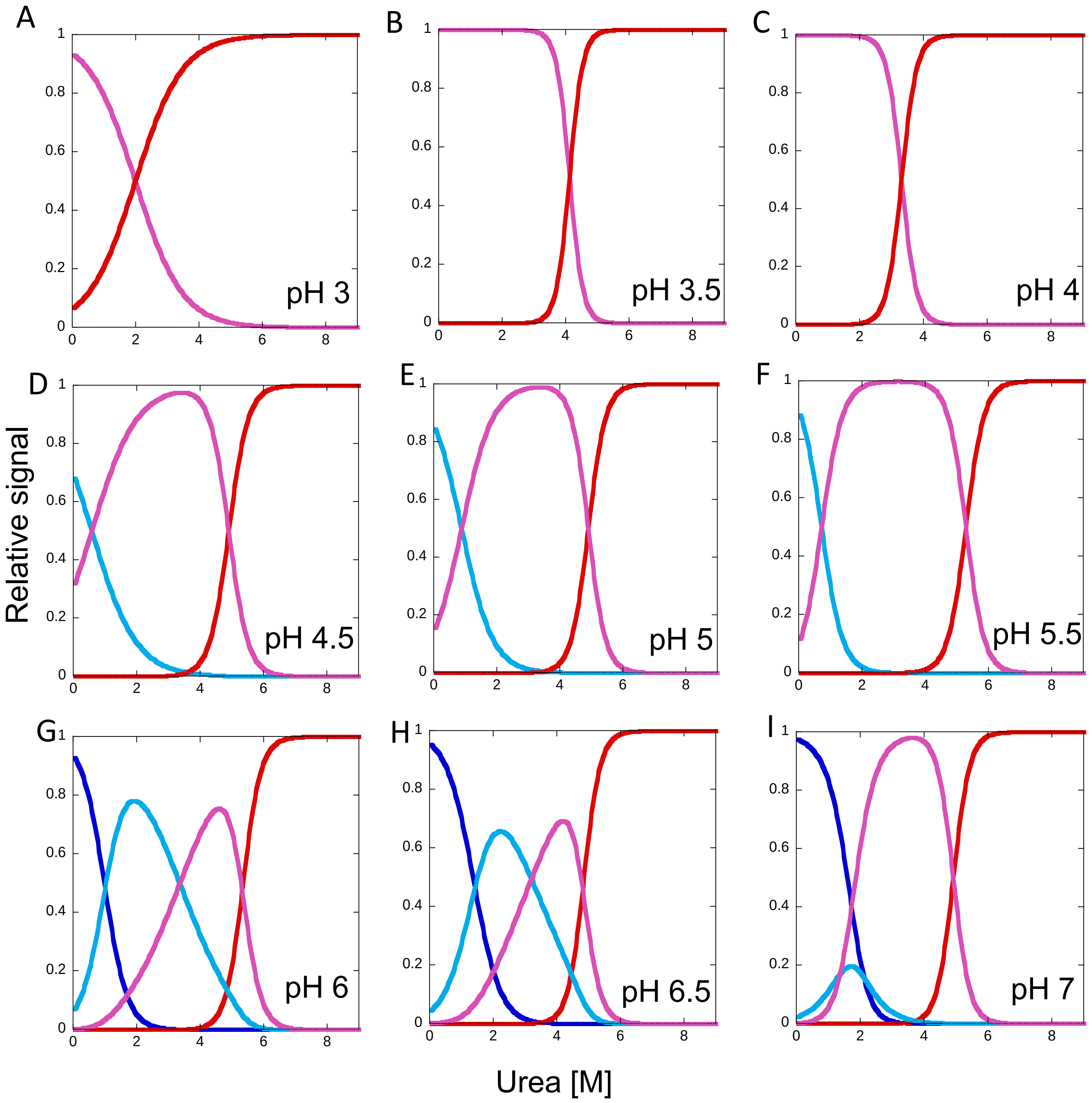

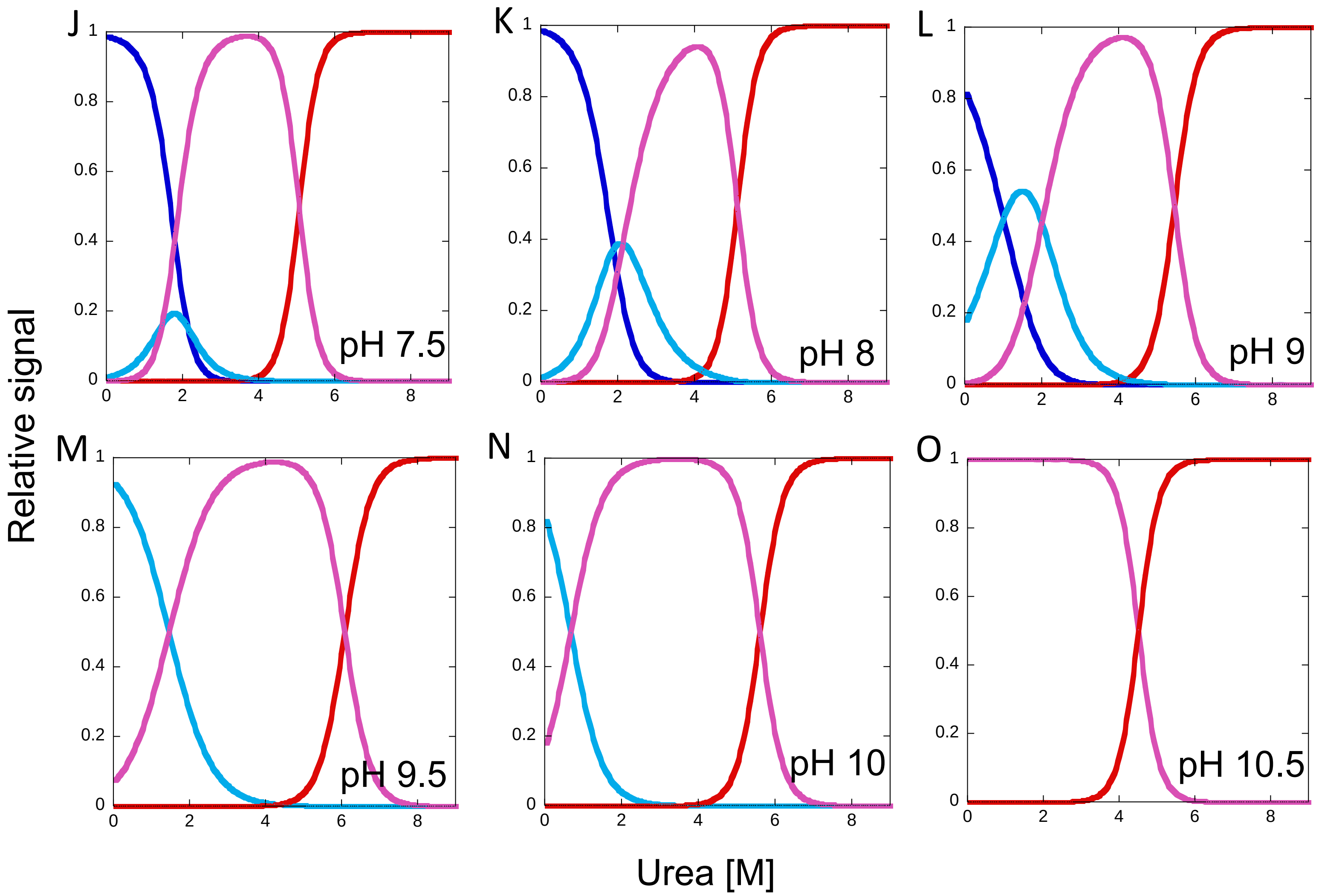


**Figure S3**.Fraction of species of PaCasp7a equilibrium unfolding as a function of urea concentration from pH 3 to pH 10.5. The fractions of native, $intermediate_{1,}$ $intermediate_{2}$ and unfolded protein were calculated as a function of urea concentration from fits through the data at each pH. Fraction of species – native (-) , $intermediate_{1}$ (-) , $intermediate_{2}$ (-) and unfolded protein (-)

**Supplementary figure S4.**

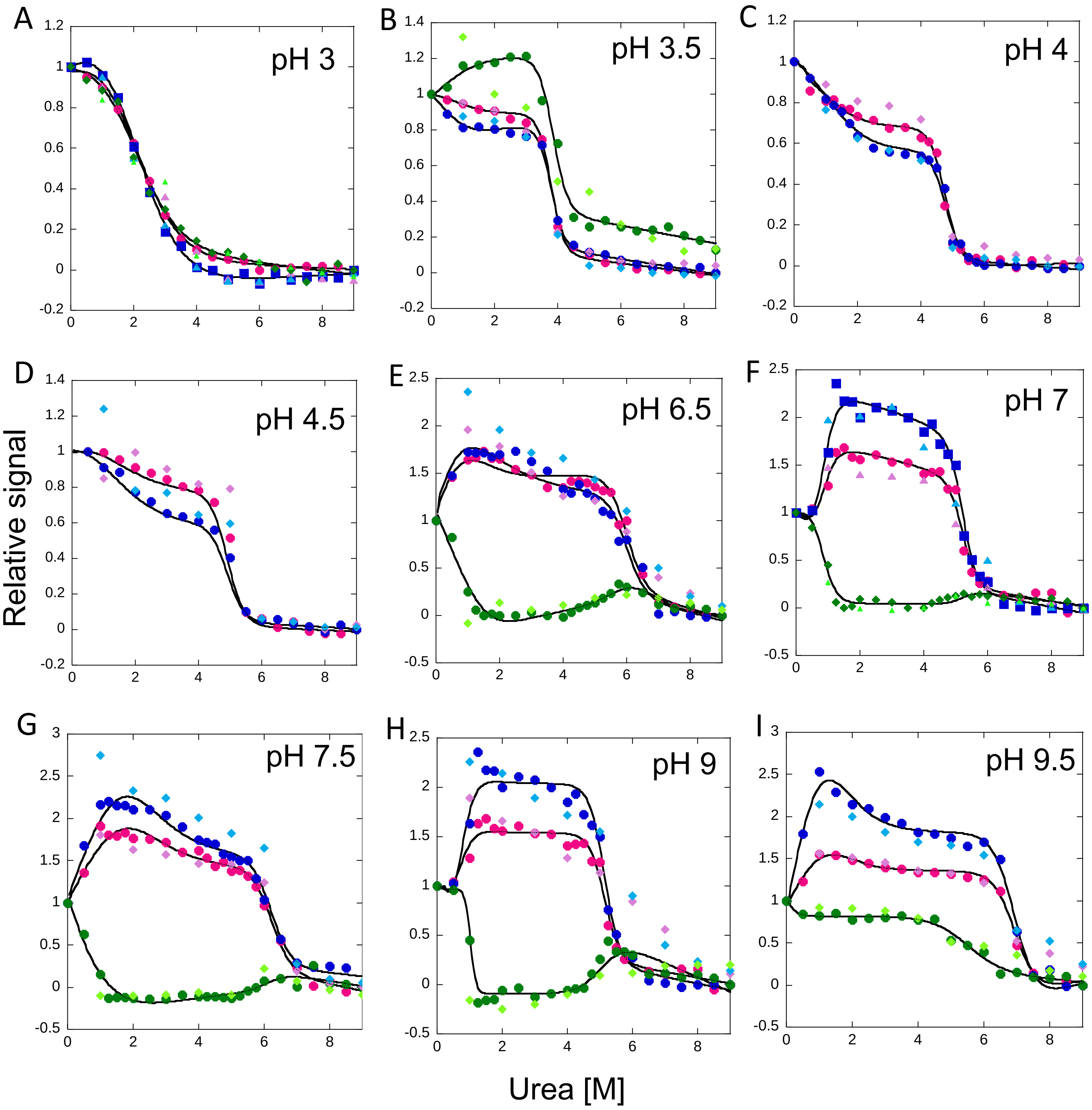

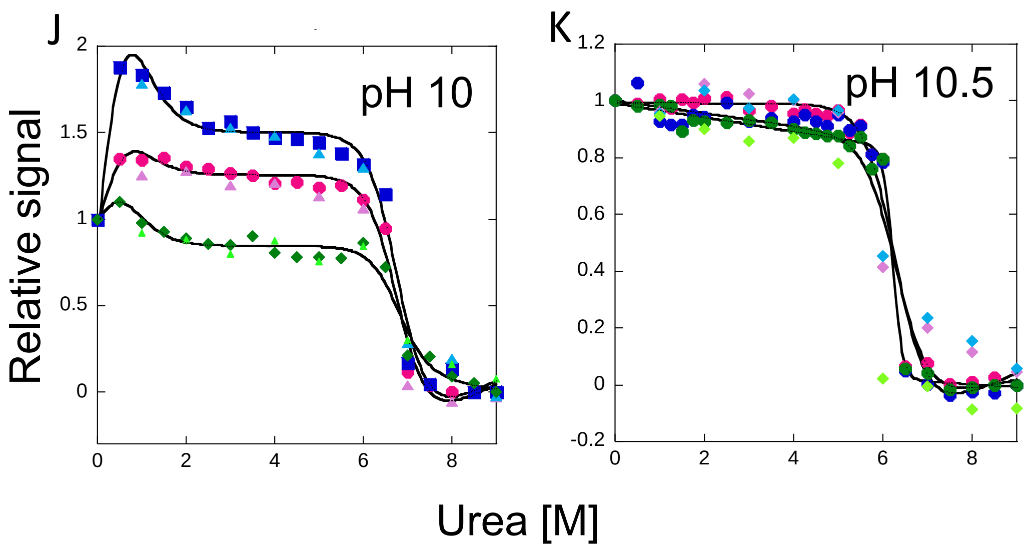


**Figure S4**. Normalized CD and equilibrium unfolding data for OfCasp3a from pH 3 to pH 10.5. Colored solid symbols represent averaged raw data and solid lines through the data represent the global fits, as described in the main text. For A-K, the following symbols were used: excitation at 280 nm- unfolding(●) and refolding(◆) , excitation at 295 nm - unfolding (●) and refolding (◆), CD data - unfolding (●) and refolding (◆)

**Supplementary figure S5.**

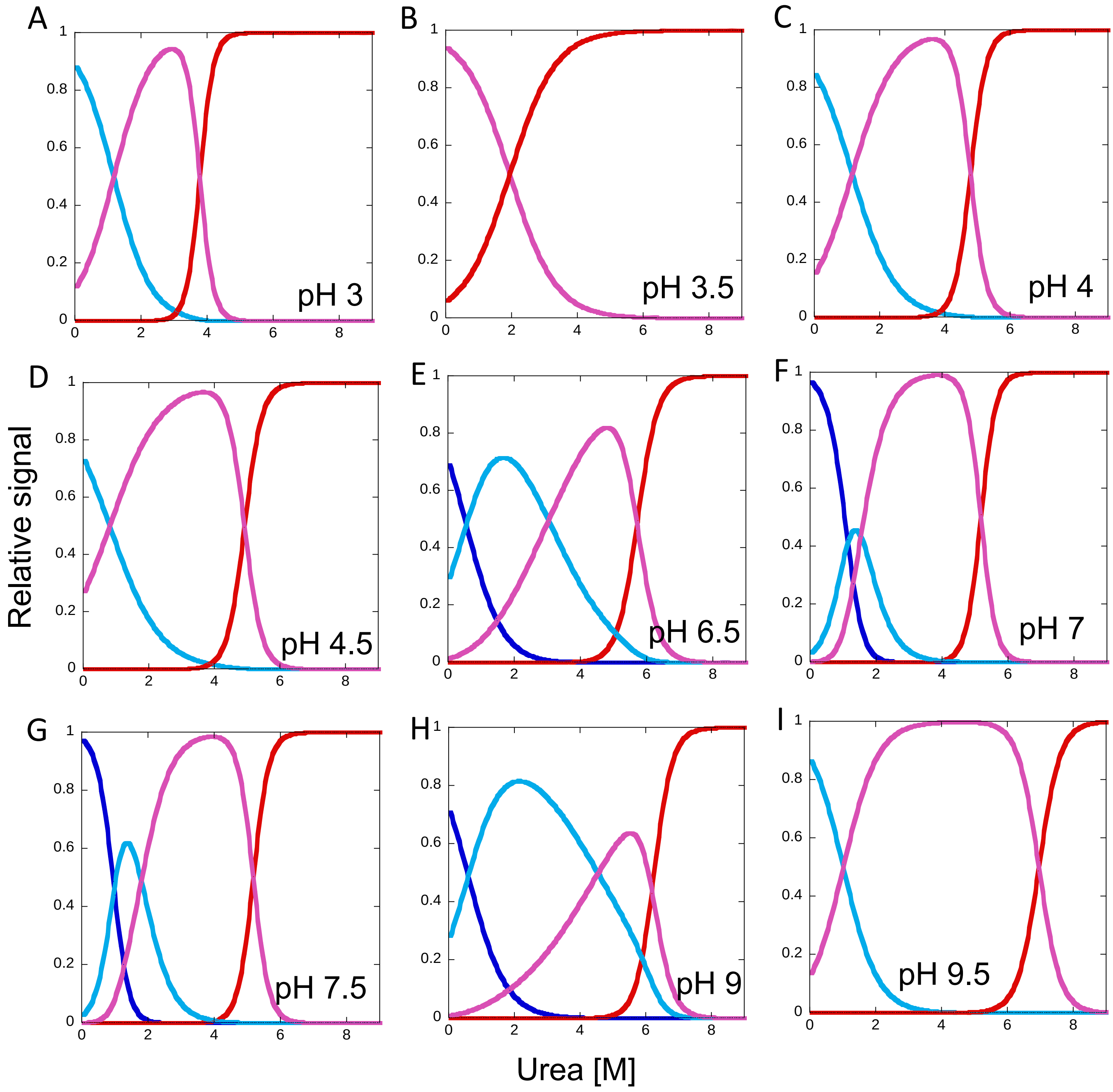

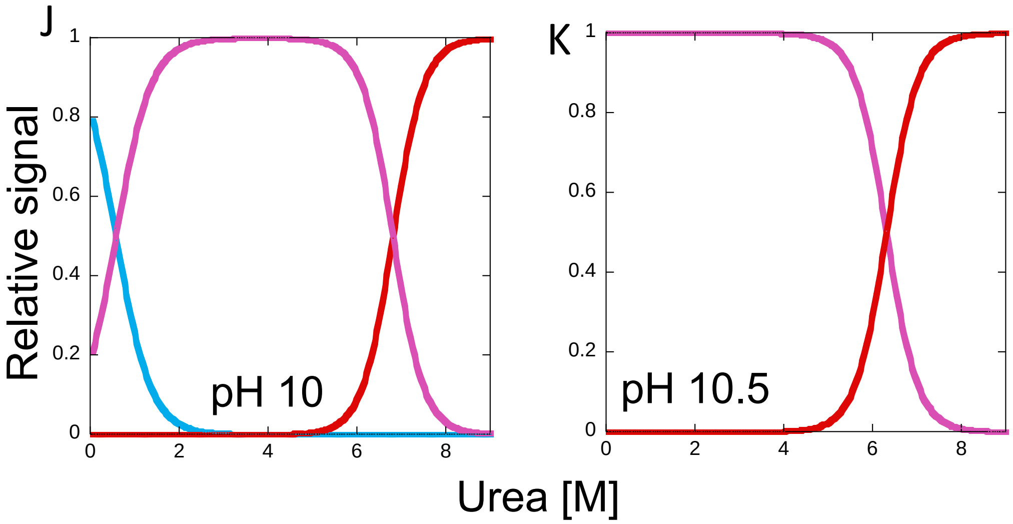


**Figure S5.** Fraction of species of OfCasp3a equilibrium unfolding as a function of urea concentration from pH 3 to pH 10.5. The fractions of native, intermediate$_{1,}$ intermediate$_2$ and unfolded protein were calculated as a function of urea concentration from fits through the data at each pH. Fraction of species - native(−) , intermediate$_1$ (−) , intermediate$_2$ (−) and unfolded protein (−)

**Supplementary table SI.** Summary of free energy changes and co-operativity index (m-values) for each transition in equilibrium unfolding of PaCasp7a

| pH | $\Delta G_1$ (kcal m $^{-1}$) | $m_1$ (kcal m $^{-1}$ M $^{-1}$) | $\Delta G_2$ (kcal m $^{-1}$) | $m_2$ (kcal m $^{-1}$ M $^{-1}$) | $\Delta G_3$ (kcal m $^{-1}$) | $m_3$ (kcal m $^{-1}$ M $^{-1}$) | Total $\Delta G°_{conf}$ (kcal mol $^{-1}$) |
|---|---|---|---|---|---|---|---|
| 3 | | | | | 1.6 ± 0.2 | 0.80 ± 0.1 | 1.6 ± 0.2 |
| 3.5 | | | | | 8.3 ± 0.6 | 2.51 ± 0.2 | 8.3 ± 0.6 |
| 4 | | | | | 11.5 ± 0.7 | 2.80 ± 0.3 | 11.5 ± 0.7 |
| 4.5 | | | 0.9 ± 0.7 | 0.89 ± 0.3 | 10.5 ± 0.6 | 2.06 ± 0.4 | 11.4 ± 0.6 |
| 5 | | | 1.0 ± 0.5 | 1.21 ± 0.4 | 10.6 ± 0.6 | 2.17 ± 0.4 | 11.6 ± 0.5 |
| 5.5 | | | 1.2 ± 0.9 | 1.78 ± 0.3 | 10.8 ± 1.0 | 2.05 ± 0.2 | 12.0 ± 0.5 |
| 6 | 1.5 ± 0.2 | 1.61 ± 0.3 | 2.2 ± 0.8 | 0.66 ± 0.3 | 10.9 ± 1.7 | 2.05 ± 0.3 | 14.6 ± 0.9 |
| 6.5 | 1.8 ± 0.2 | 1.30 ± 0.2 | 2.0 ± 0.4 | 0.63 ± 0.1 | 10.6 ± 0.9 | 2.21 ± 0.4 | 14.4 ± 0.9 |
| 7 | 2.2 ± 0.3 | 1.05 ± 0.3 | 1.5 ± 0.4 | 1.10 ± 0.1 | 11.2 ± 0.1 | 2.30 ± 0.3 | 14.9 ± 0.5 |
| 7.5 | 2.6 ± 0.9 | 1.23 ± 0.3 | 1.8 ± 0.9 | 1.30 ± 0.3 | 11.9 ± 1.2 | 2.32 ± 0.3 | 16.3 ± 0.3 |
| 8 | 2.5 ± 0.9 | 1.30 ± 0.3 | 2.0 ± 1.0 | 0.95 ± 0.1 | 11.5 ± 1.0 | 2.21 ± 0.1 | 16.0 ± 1.0 |
| 9 | 0.9 ± 0.8 | 0.93 ± 0.3 | 2.2 ± 0.5 | 1.12 ± 0.2 | 10.9 ± 0.1 | 2.05 ± 0.3 | 13.9 ± 0.9 |
| 9.5 | | | 1.5 ± 0.4 | 1.05 ± 0.1 | 10.6 ± 0.8 | 1.75 ± 0.2 | 12.1 ± 0.3 |
| 10 | | | 0.9 ± 0.3 | 1.41 ± 0.3 | 11.0 ± 0.1 | 1.97 ± 0.2 | 11.9 ± 0.6 |
| 10.5 | | | | | 9.7 ± 0.9 | 2.18 ± 0.3 | 9.7 ± 0.2 |

**Supplementary table SII.** Summary of free energy changes and co-operativity index (m-values) for each transition in equilibrium unfolding of OfCasp3a

| **pH** | **$\Delta G_1$ (kcal m $^{-1}$)** | **$m_1$ (kcal m $^{-1}$ M $^{-1}$)** | **$\Delta G_2$ (kcal m $^{-1}$)** | **$m_2$ (kcal m $^{-1}$ M $^{-1}$)** | **$\Delta G_3$ (kcal m $^{-1}$)** | **$m_3$ (kcal m $^{-1}$ M $^{-1}$)** | **Total $\Delta G°_{conf}$ (kcal mol $^{-1}$)** |
|---|---|---|---|---|---|---|---|
| 3 | | | | | 1.7 ± 0.3 | 0.86 ± 0.1 | 1.7 ± 0.3 |
| 3.5 | | | 1.1 ± 0.5 | 1.05 ± 0.3 | 10.8 ± 0.6 | 2.89 ± 0.2 | 12.0 ± 0.5 |
| 4 | | | 1.0 ± 0.4 | 0.90 ± 0.2 | 11.5 ± 0.5 | 2.41 ± 0.1 | 12.5 ± 0.4 |
| 4.5 | | | 1.0 ± 0.5 | 0.81 ± 0.3 | 12.1 ± 0.6 | 2.36 ± 0.3 | 13.1 ± 0.5 |
| 6.5 | 0.8 ± 0.2 | 0.44 ± 0.1 | 1.8 ± 0.5 | 0.60 ± 0.2 | 12.5 ± 0.9 | 2.09 ± 0.2 | 15.1 ± 0.5 |
| 7 | 2.0 ± 0.4 | 1.80 ± 0.3 | 1.9 ± 0.5 | 1.20 ± 0.5 | 13.4 ± 0.6 | 2.60 ± 0.8 | 17.3 ± 0.5 |
| 7.5 | 2.1 ± 0.5 | 2.20 ± 0.4 | 2.2 ± 0.4 | 1.28 ± 0.3 | 12.9 ± 0.6 | 2.50 ± 0.3 | 17.2 ± 0.5 |
| 9 | 0.5 ± 0.4 | 1.0 ± 0.3 | 2.0 ± 0.8 | 0.45 ± 0.3 | 12.5 ± 2.4 | 2.03 ± 0.1 | 15 ± 0.3 |
| 9.5 | | | 1.0 ± 0.3 | 1.13 ± 0.1 | 12.7 ± 0.7 | 1.83 ± 0.1 | 13.7 ± 0.5 |
| 10 | | | 0.8 ± 0.2 | 1.50 ± 0.3 | 11.5 ± 0.1 | 1.70 ± 0.2 | 12.3 ± 0.1 |
| 10.5 | | | | | 10.5 ± 0.2 | 1.67 ± 0.2 | 10.5 ± 0.2 |

**Supplementary figure S6.**

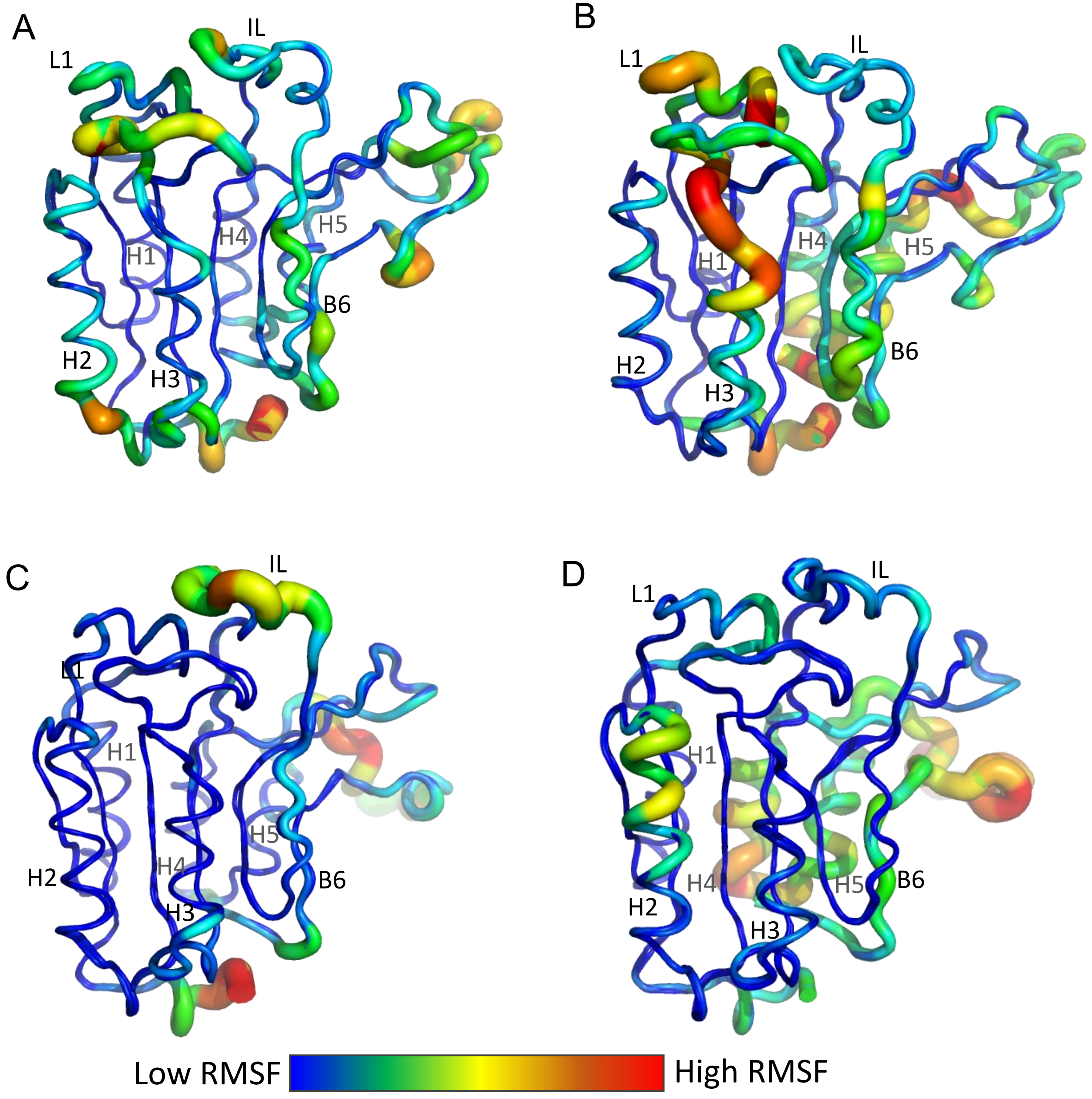


**Figure S6.** RMSF values from MD simulations converted to B-factors, overlaid and mapped onto the modeled structure of PaCasp7a (top panels) and OfCasp3a (bottom panels). PaCasp7a in water (A) or in 5M urea(B); OfCasp3a in water (C) or in 5M urea (D).

**Supplementary figure S7.**

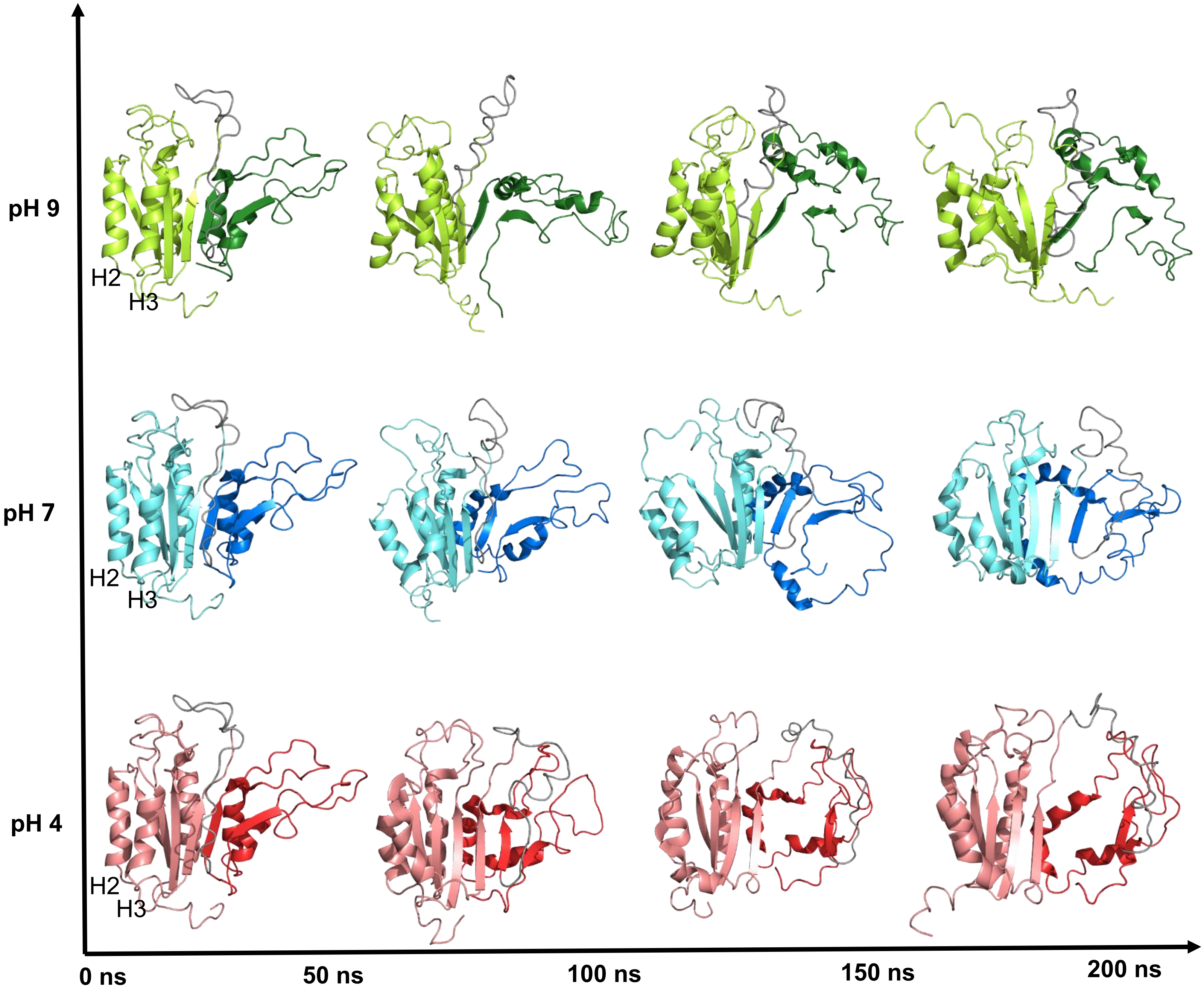


**Figure S7**. Urea MD snapshots of PaCasp7a at pH 4 (LS - salmon, SS - red), pH 7 (LS - cyan, SS - blue), and pH 9 (LS - limon, SS - green) with helices 2 and 3 (H2, H3) labeled on the structure at 0 ns.

**Supplementary figure S8.**

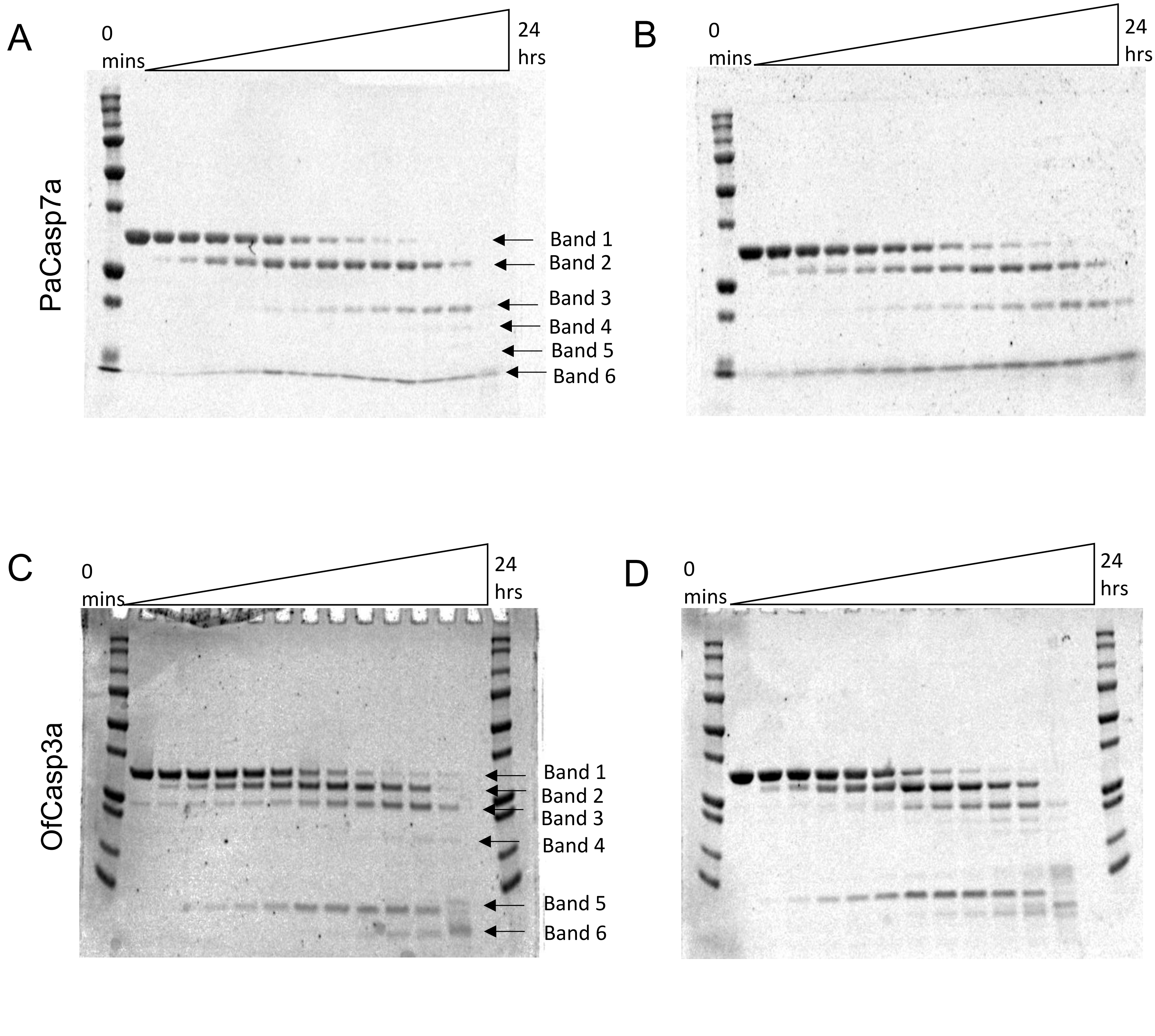


**Figure S8.** Limited trypsin digestion of PaCasp7a (A,B) and of OfCasp3a (C,D) at pH 7 (panels B and D) and pH 9 (panels C and D). The 31 kDa band observed at 0 min represents the native proteins, whereas the 26 kDa (band 2), 20 kDa (band 3), 18 kDa (band 4), 15 kDa (band 5), and 9 kDa (band 6) are the cleavage products as described in the text.

**Supplementary table SIII.** Caspases used in the phylogenetic analysis and their respective accession numbers.

| **Caspases** | **Accession number** |
|---|---|
| *Homo sapiens* | |
| HsCasp-2 | NP_116764.2 |
| HsCasp-3 | NP_004337.2 |
| HsCasp-4 | NP_001216.1 |
| HsCasp-5 | NP_004338.3 |
| HsCasp-6 | NP_001217.2 |
| HsCasp-7 | NP_001253985.1 |
| HsCasp-8 | NP_001219.2 |
| HsCasp-9 | NP_001220.2 |
| HsCasp-10 | NP_116759.2 |
| *Mus musculus* | |
| MmCasp-2 | NP_031636.1 |
| MmCasp-3 | NP_001271338.1 |
| MmCasp-6 | NP_033941.3 |
| MmCasp-7 | XP_006526679.1 |
| MmCasp-8 | NP_001264855.1 |
| MmCasp-9 | NP_056548. |
| *Gallus gallus* | |
| GgCasp-2 | NP_001161173.1 |
| GgCasp-3 | NP_990056.1 |
| GgCasp-6 | NP_990057.1 |
| GgCasp-7 | XP_421764.3 |
| GgCasp-8 | NP_989923.1 |
| GgCasp-9 | XP_424580.5 |
| GgCasp-10 | XP_421936.4 |
| *Xenopus tropicalis* | |
| XtCasp-2 | XP_012809163.2 |
| XtCasp-3 | NP_001120900.1 |
| XtCasp-6 | NP_001011068.1 |
| XtCasp-7 | NP_001016299.1 |
| XtCasp-8 | XP_017953067.2 |
| XtCasp-9 | NP_001116935.1 |
| XtCasp-10 | NP_001015715.2 |

| Caspases | Accession number |
|---|---|
| *Danio rerio* | |
| DrCasp-2 | NP_001036160.1 |
| DrCasp-3a | XP_001338890.2 |
| DrCasp-3b | XP_005173133.1 |
| DrCasp-6a | XP_005164109.1 |
| DrCasp-6b | XP_017210076.1 |
| DrCasp-6c | NP_001018333.1 |
| DrCasp-7 | XP_005156389.1 |
| DrCasp-8 | NP_001092089.1 |
| DrCasp-9 | NP_001007405.2 |
| *Coral caspases* | |
| Acropora digitifera Casp-3 | XP_015775441.1 |
| Acropora digitifera Casp-8 | XP_015761120.1 |
| Hydra vulgaris Casp-2 | NP_001274285.1 |
| Hydra vulgaris Casp-3 | XP_012557085.1 |
| Hydra vulgaris Casp-8 | XP_012562456.1 |
| Orbicella faveolata Casp-3a | XP_020613409.1 |
| Orbicella faveolata Casp-3b | XP_020630525.1 |
